# Robust biomarker discovery through multiplatform multiplex image analysis of breast cancer clinical cohorts

**DOI:** 10.1101/2023.01.31.525753

**Authors:** Jennifer Eng, Elmar Bucher, Zhi Hu, Melinda Sanders, Bapsi Chakravarthy, Paula Gonzalez, Jennifer A. Pietenpol, Summer L. Gibbs, Rosalie C. Sears, Koei Chin

## Abstract

Spatial profiling of tissues promises to elucidate tumor-microenvironment interactions and enable development of spatial biomarkers to predict patient response to immunotherapy and other therapeutics. However, spatial biomarker discovery is often carried out on a single patient cohort or imaging technology, limiting statistical power and increasing the likelihood of technical artifacts. In order to analyze multiple patient cohorts profiled on different platforms, we developed methods for comparative data analysis from three disparate multiplex imaging technologies: 1) cyclic immunofluorescence data we generated from 102 breast cancer patients with clinical follow-up, in addition to publicly available 2) imaging mass cytometry and 3) multiplex ion-beam imaging data. We demonstrate similar single-cell phenotyping results across breast cancer patient cohorts imaged with these three technologies and identify cellular abundance and proximity-based biomarkers with prognostic value across platforms. In multiple platforms, we identified lymphocyte infiltration as independently associated with longer survival in triple negative and high-proliferation breast tumors. Then, a comparison of nine spatial analysis methods revealed robust spatial biomarkers. In estrogen receptor-positive disease, quiescent stromal cells close to tumor were more abundant in good prognosis tumors while tumor neighborhoods of mixed fibroblast phenotypes were enriched in poor prognosis tumors. In triple-negative breast cancer (TNBC), macrophage proximity to tumor and B cell proximity to T cells were greater in good prognosis tumors, while tumor neighborhoods of vimentin-positive fibroblasts were enriched in poor prognosis tumors. We also tested previously published spatial biomarkers in our ensemble cohort, reproducing the positive prognostic value of isolated lymphocytes and lymphocyte occupancy and failing to reproduce the prognostic value of tumor-immune mixing score in TNBC. In conclusion, we demonstrate assembly of larger clinical cohorts from diverse platforms to aid in prognostic spatial biomarker identification and validation.

**Statement of significance:** Our single-cell spatial analysis of multiple clinical cohorts uncovered novel biomarkers of patient outcome in breast cancer. Additionally, our data, software, and methods will help advance spatial characterization of the tumor microenvironment.

## Introduction

Recent advances in the treatment landscape of breast cancer have motivated the characterization of the breast tumor microenvironment for deeper understanding of immune-tumor interactions. Identification of biomarkers predicting breast cancer immunotherapy response is still an urgent clinical need^1^. For instance, in metastatic TNBC, only a quarter of PD-L1 positive patients respond to single-agent immune checkpoint blockade^1^. In contrast, in early stage TNBC, response rates to neoadjuvant immune checkpoint plus chemotherapy were similar in PD-L1 positive and negative groups^2^. This highlights the need for biomarker development for better patient stratification across disease stages. Furthermore, trials evaluating novel immune targeted therapies in breast cancer should be accompanied by biomarker development for maximum therapeutic efficacy^3^.

Highly multiplex imaging methods enable quantification of dozens of biomarkers in a single tissue section with at sub-cellular resolution while retaining spatial context^4–9^. Tissue structures such as tertiary lymphoid structures, identified with multiplex imaging, are predictive biomarkers of immunotherapy response in melanoma^10,11^. Spatial proximity between tumor, immune and stromal cell types is associated with response to neoadjuvant therapy in HER2+ breast cancer^12^. In several breast cancer multiplex imaging studies, single-cell spatial context has prognostic relevance and shows correlations with transcriptomic and genomic features of tumors^13–16^. However, spatial biomarkers can be difficult to reproduce due to limited numbers of patients used to develop them and difficulties in comparing data from different imaging platforms. Furthermore, overfitting is an issue in biomedical imaging data due to the number of steps in the processing pipeline and the number of variables and parameters involved. Overfitting can be addressed through use of a discovery cohort to tune analytical methods, which are then fixed and subsequently applied to a validation cohort^17^. In theory, validation cohorts can be readily obtained through incorporation of publicly available data from disparate imaging platforms into biomarker studies. In practice, integrated analysis of such data remains a challenge. Furthermore, documentation of metadata, analysis protocols and code is essential for reuse of data and reproducibility of findings, preferably using open-source software tools^17^. We developed an open-source python software, mplexable^18^, for multiplex image processing and analysis, which we use herein to process and analyze three multiplex imaging breast cancer cohorts: a cyclic immunofluorescence (CyCIF) dataset which we generated, and publicly available imaging mass cytometry (IMC) and multiplex ion-beam imaging (MIBI) datasets^13,15,19^. In this proof-of-concept study, we identify prognostic single-cell spatial biomarkers common across the imaging platforms. As such imaging datasets become more widely available, tools such as ours can facilitate biomarker discovery with high accuracy, reliability, and efficiency.

## Methods

### Patient samples

Two breast cancer tissue microarrays were graciously provided by Dr. Jennifer Pietenpol (JP). All samples were collected at time of surgical resection (mastectomy or breast conserving surgery) at Vanderbilt University Medical Center with the same fixation protocol. JP-TMA1 had 131 cores of approximately 1.2 μm diameter, with duplicate cores from 19 TNBC, 8 HER2+ and 36 ER+ patients. It also included one control core each of inflamed appendix, colon cancer, muscle, pancreas and normal breast. Four of the TMA1 TNBC patients received neoadjuvant chemotherapy. JP-TMA2 contained a single, slightly larger (∼1.4μm diameter) core from 39 triple-negative tumors and 1 ER+/HER2+ control core. Thirteen of the patients in TMA2 received neoadjuvant therapy. Clinical outcome and clinicopathological information are available for TMA1 and TMA2.

### Imaging data generation and sources

CyCIF staining of tumor tissue was completed on JP-TMA1 and TMA2. The whole tissue core was imaged, as described previously^18^. MIBI imaging data was previously published by Keren *et al*^13^ and the images were downloaded from https://mibi-share.ionpath.com/tracker/imageset under the name “Keren et al., Triple Negative Breast Cancer.” Survival and recurrence data were obtained from a second publication by the same group^20^, and were downloaded from https://github.com/aalokpatwa/rasp-mibi. IMC imaging data were previously published by Jackson *et al*.^15^ and images and clinical data were downloaded from https://doi.org/10.5281/zenodo.3518284.

### Image Processing

CyCIF tiff images were registered, segmented and single-cell intensity as well as nuclear size and shape features were extracted as previously described^18^. The CyCIF pipeline is available at https://gitlab.com/engje/mplexable/-/tree/master/jupyter. Nuclear and cell segmentation were run using the Cellpose algorithm^21^, which showed visually superior performance on CyCIF data compared to a watershed algorithm (S1a). Nuclear and cell segmentation masks were matched using mplexable, enabling subtraction of nuclear mask from cell mask to obtain segmentation of the cytoplasm (S1b).

MIBI and IMC processing pipelines documenting the steps described here are available as Jupyter notebooks at https://github.com/engjen/cycIF_TMAs. MIBI and IMC images were downloaded as multipage OMEtiffs. Hot pixels^14^ were detected by identifying pixels that were 10 standard deviations above a median filtered image with a 2×2 pixel kernel size. Hot pixels were set to the median filter values and resulting images were saved as tiffs for downstream feature extraction. DNA channels were processed for nuclear segmentation as follows. DNA images were rescaled between the 3^rd^ and one and a half times the 99.999 quantile. The gamma value was adjusted by 0.6 in MIBI data and 0.4 in IMC data to enhance dimly stained nuclei. A two-channel nuclear plus cytoplasm image was generated for cell segmentation. A mixture of markers was used to generate a composite image which stained the cytoplasm of the majority of cells. For MIBI data, the β-catenin, vimentin, CD45 and CD31 channels were combined into a maximum intensity projection cytoplasm image and the gamma value was adjusted by 0.6. For IMC data, E-cadherin, vimentin, CD44 and CD45 were combined into a maximum intensity projection cytoplasm image and gamma adjusted by 0.4. Chamboelle total variation denoising, implemented in scipy^22^, was used to smooth out pixelated nuclear and cytoplasmic projection images (weight=0.1, except weight=0.05 for IMC cytoplasm). All parameters were selected by testing segmentation results at https://www.deepcell.org/predict and <u>cellpose.org</u> (S1c-d). Skipping either nuclear or cytoplasmic Chamboelle total-variation de-noising resulted in failure of deep learning-based algorithms on the IMC data (S1e). Mesmer segmentation^23^ performed better than Cellpose^24^ in IMC data due to improved detection of dim nuclei, likely due to the incorporation of cytoplasmic staining in the nuclear segmentation model (S1d). This issue with Cellpose was not detected in CyCIF images, which had brighter DNA staining. For IMC and MIBI data, nuclear and cellular segmentation were performed on processed nuclear and nuclear + cytoplasm images using Mesmer^23^. Matching of cell IDs in the nuclear and cell masks was done with mplexable^18^, with cell masks relabeled to match the ID of the nucleus which they had most overlap with. Cytoplasm masks were calculated by subtracting the nuclear mask from the matching cell mask. Nuclear and cytoplasmic mean intensity, nuclear size and shape features, and nuclear centroid coordinates were extracted with mplexable^18^.

Mesmer, a deep-learning-based approach for nuclear and whole-cell segmentation of tissue data, was run on IMC data and compared to the watershed-based segmentation originally published by Jackson *et al*^15^. The cell counts across the two methods had a Pearson correlation of 0.98 (S2a). Visual examination of ROIs with discordant cell numbers revealed that Mesmer segmentation performed better in tissues with necrosis and high background noise in the DNA channel (S2 b-d).

### Image QC

In IMC data, artifacts include non-specific background staining, necrotic regions, and bright antibody aggregates. IMC data were collected from small ROIs (∼600 μm) within TMA cores and some samples annotated as estrogen-receptor (ER)+ tissues did not show any ER+ staining in the ROI. Therefore, quality control (QC) was performed on ER-stained images, a marker which has been noted to exhibit non-specific background staining on the IMC platform^25^. QC images of ER staining were generated and sorted in a blinded fashion into negative and positive for nuclear-specific-staining (S3a). ROIs from clinically annotated ER+ patients that were classified as ER positive during QC or ROIs that came from ER negative patients and classified as ER negative were used for analysis. Samples that passed ER QC did not have significantly different grade, PR status, TMA block, age of specimen, age of patient or tumor size (S3c-d). There were no significant survival differences between QC passed versus failed tumors from ER+, TNBC or ER+HER2+ patients (S3e). In the IMC dataset, additional QC steps included: necrotic regions were manually circled using the napari^26^ image viewer and excluded and bright aggregates in the CD3 channel were excluded by removing cells above a threshold set at the value of CD3+ cells showing an appropriate membranous staining pattern.

In the CyCIF data, imaging artifacts included autofluorescence (AF), non-specific background, floating tissue and tissue loss. Background AF images were obtained half-way through CyCIF data collection and these images were scaled by exposure time and subtracted from the AF488, AF55 and AF647 channels using mplexable^18^. Feature extraction was performed on AF subtracted images. Areas of floating tissue, air bubbles or necrotic regions were manually circled using the napari^26^ image viewer and excluded. Non-specific background staining was removed by setting manual thresholds for selected markers and subtracting those values from extracted data. The PD1 antibody had bright aggregates that were excluded with an upper threshold. Tissue loss was detected by cells that lacked DAPI staining in the last round of imaging, and these cells were excluded.

In all three platforms, additional artifacts caused by floating tissue or imaging problems (e.g., dark or bright bands across IMC and MIBI images perhaps caused by problems with the rastering process) were detected through unsupervised clustering and visual inspection of clusters on the images. Clusters comprised of artifacts showed atypical very bright or dim staining in many channels, formed distinct artifact clusters and were removed.

### Single cell phenotyping

Cell types were defined in two ways: manual gating and unsupervised clustering. Unsupervised clustering was conducted using the scanpy^27^ software. Single-cell mean intensity values were selected from either the nucleus or cytoplasm masks for each marker, depending on expected intracellular distribution. β-catenin, which can localize on the membrane, in the cytoplasm or in the nucleus, was selected from both nuclear and cytoplasmic masks for separate analysis. Since the CyCIF and IMC platforms had more marker and breast cancer subtype overlap than the MIBI panel (**Figure 1**, S4) 20 matching markers were selected for clustering in these datasets, plus selected markers for immune, epithelial and fibroblast subsets (collagen I [ColI], CD4, CD8 in CyCIF, and fibronectin [FN], pan-cytokeratin [panCK] in IMC). For MIBI data, all available markers were used for clustering. Additionally, the nuclear area feature was used for clustering. Each marker was divided by its standard deviation, without zero-centering, and clipped above 20 standard deviations. A Uniform Manifold Approximation and Projection (UMAP) embedding was generated using 30 k-nearest neighbors and the embedding was clustered using the Leiden community detection algorithm^28^. The Leiden resolution parameter was selected that resulted in 20 - 25 clusters. Each cluster was annotated and categorized as epithelial, endothelial, fibroblast, immune or stromal. Some clusters were comprised of multiple cell types, and these were manually split, for example, the CD44+ cluster was split into CD44+ tumor and CD44+ stroma based on manual gating results (described below).

**Figure 1.**
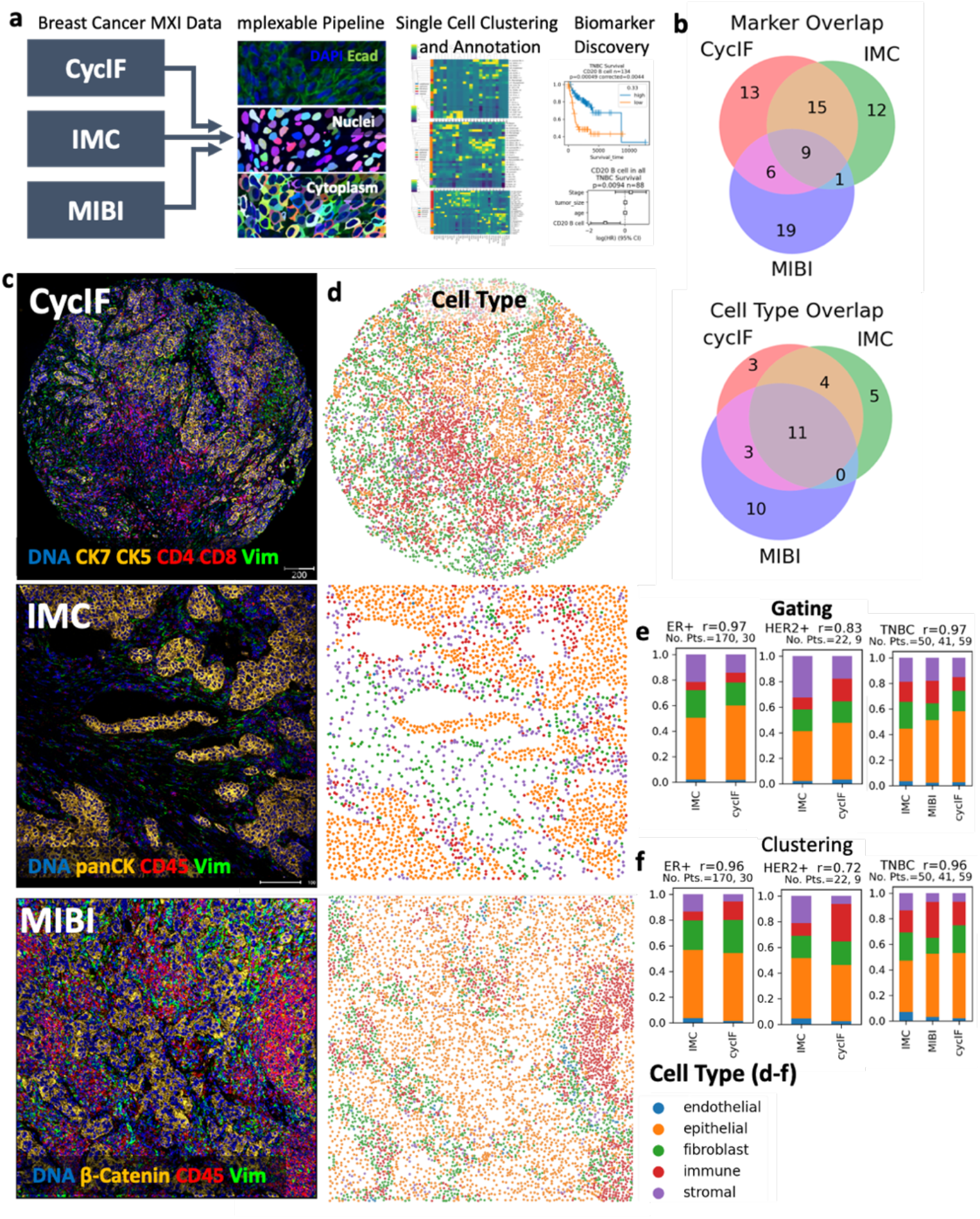
Analysis of breast cancer multiplex imaging datasets from different platforms. a. Three multiplex imaging (MXI) datasets from breast cancer tissue microarrays were processed through single cell segmentation and feature extraction using the mplexable pipeline. The single cell datasets were separately clustered using the unsupervised Leiden algorithm resulting in cell types which were annotated with similar names across platforms. Finally, the data were used for discovery and validation of prognostic cell abundance and spatial biomarkers. cycIF, cyclic immunofluorescence; IMC, imaging mass cytometry; MIBI, multiplex ion beam imaging. b. Overlap of markers (top) and annotated cell types (bottom) in each multiplex imaging dataset. c. Representative images from the three multiplex imaging platforms showing epithelial (orange), immune (red) and fibroblast (green) markers. d. Gated cell types showing cell location and lineages: epithelial (orange), immune (red), fibroblast (green), endothelial (blue) and other stromal (purple). e-f. Cell lineage fraction of total cells per subtype, per platform using either manual gating (e) or unsupervised clustering and annotation (f) to determine lineage. Pearson’s correlation between platforms (r=0.xx) and number of patients for each subtype and platform shown in figure title.

We then performed manual gating to verify our annotated-cluster cell type. A threshold was set for each gating marker based on the expected morphology of positive staining in images. Fibroblasts were defined as positive for one or more of vimentin, fibronectin (FN) or collagen I (ColI). Epithelial cells were defined as positive for one or more of Ecad, cytokeratins, or β-catenin. Endothelial cells were defined as CD31+. Immune cells were defined as CD45+. Stromal cells were defined as all non-fibroblast, non-endothelial, non-epithelial, non-immune segmented nuclei. Jupyter notebooks recording single cell phenotyping pipelines are available here: https://github.com/engjen/cycIF_TMAs

### Single Cell Based Patient Subtyping

Epithelial and stromal subtypes were determined by unsupervised clustering of patients based on the fraction of epithelial or stromal cell types within each compartment, respectively. Cell types representing greater than 2-4% of the total cell population in the respective tissue compartment were used for clustering. This cutoff was chosen to ignore rare, platform specific cell types that may represent method-specific artifacts. Unsupervised clustering of patients was performed using the Leiden algoritm implemented in scanpy^27^. For epithelial subtypes, the resolution of clustering was selected to minimize differences between the platforms (**Figure 2**). For stromal subtypes, we selected the minimum number of clusters need to separate T cells from other clusters (k=6).

**Figure 2.**
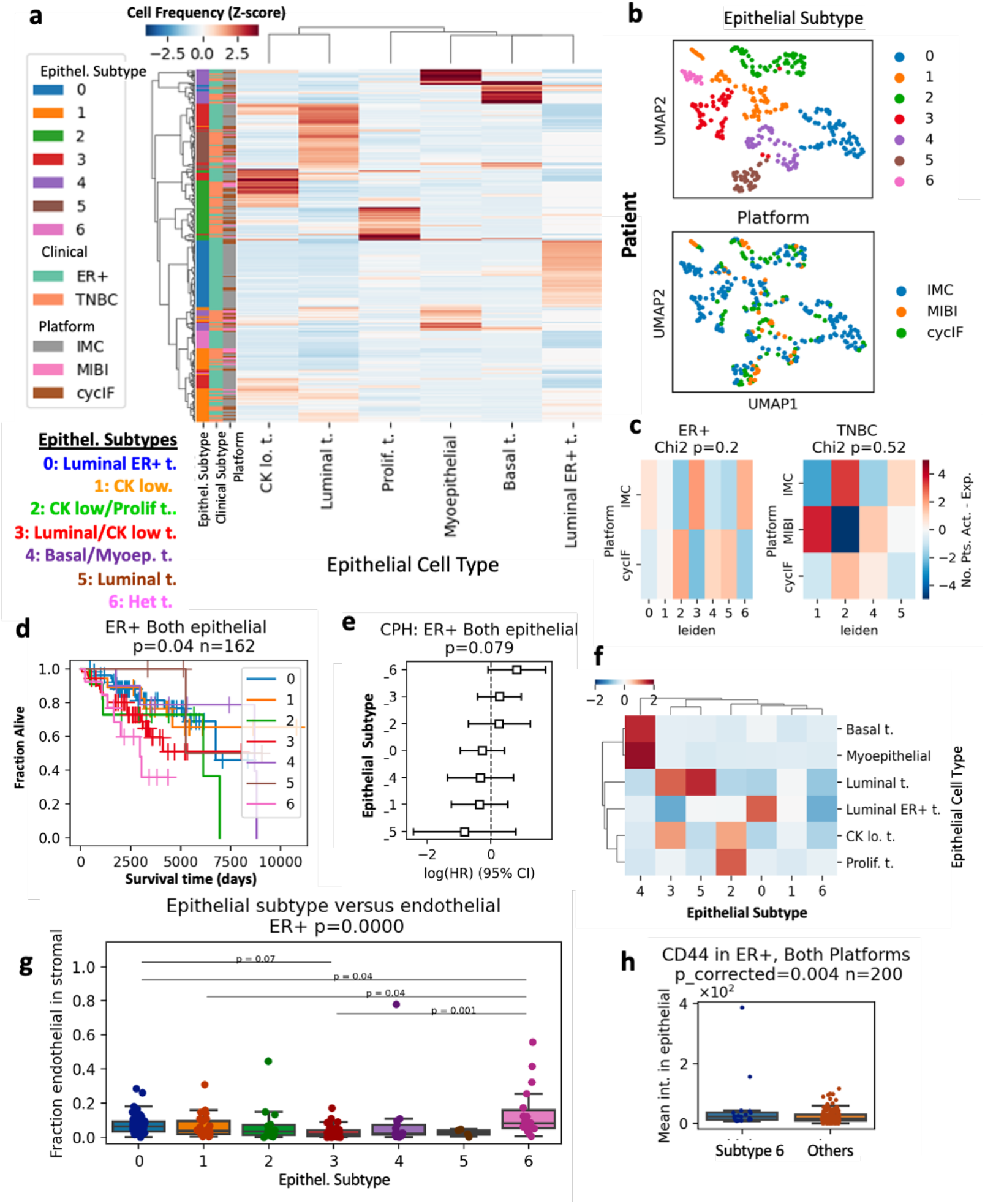
Prognostic breast cancer subtypes in combined multiplex imaging data. a. All ER+ and TNBC patients were clustered based on the fraction in each patient’s tissue of the six most common epithelial cell types, resulting in seven epithelial (Epithel.) subtypes. b. UMAP embedding of patients by fraction of epithelial cell types in all tumor cells, colored by epithelial subtype (top) and platform (bottom). c. Chi-squared analysis of epithelial subtypes versus platform, p-values in figure title. d. Kaplan-Meier curves (p-value from log-rank test) comparing overall survival (OS) in the seven epithelial subtypes present in ER+ tumors. N number of patients given in figure title. e. Cox proportional hazard (CPH) model estimating hazard ratios for epithelial subtypes of ER+ tumors. f. Mean abundance of epithelial cell types per subtype. g. Fraction of endothelial cells in tissue stromal cells of each epithelial subtype. Kruskal-Wallis H-test P-value given in figure title. Post-hoc Tukey HSD used for pairwise comparisons between groups. h. CD44 intensity in epithelial cells from poor prognosis epithelial subtype 6 compared to other ER+ patients. P-value FDR corrected t-test test given in figure title.

### Survival Analysis

For each cell type, fraction of cells of that type over all cells from combined ROIs/cores from each patient was calculated and binarized into high and low abundance based on tertile or median values within each subtype. The CyCIF TMA1 dataset was used as a discovery dataset to determine the quantile separating high and low abundance of each cell type or spatial metric that was most predictive of survival. Three quantiles were tested: 0.33 (e.g. split patients into 1/3 low and 2/3 high), 0.5 and 0.66. The most prognostic cutoff value was selected for each cell type and for cell types having prognostic value (alpha<0.05) these cutoffs were applied in the validation dataset which included CyCIF TMA2, MIBI and IMC samples. Since overall cell type fractions differed between platforms and subtypes (**Figure 1**), high and low values were determined relative to other samples from the same platform and subtype, using the predetermined cutoffs. In the validation cohort, the log-rank test p-values were corrected for multiple testing using the Benjamini– Hochberg False Discovery Rate (FDR). For biomarkers with FDR < 0.1, multivariate Cox proportional hazards (CPH) modelling was used to combine imaging biomarkers with patient age, tumor size and clinical stage to test if they were independently prognostic. Collectively, 89 TNBC and 160 ER+ patients had these additional clinical parameters (additional clinical parameters were not available in the MIBI dataset).

### Spatial Analysis

Spatial distributions of cells were calculated in multiple ways. For analysis of cellular neighbors^12^ and homotypic/heterotypic interactions^14^ each cell’s neighbors within a 40 μm radius were counted. For tumor-immune mixing score a 25 μm radius was selected to replicate Keren *et al*.^13^ For lymphocyte clusters, a 20 μm radius was used and for lymphocyte occupancy a 50 μm grid square was used, both to replicate Wortman *et al*.^16^ Ripley’s L (a density-normalized measure of clustering) and the multitype K function (Kcross; a density-normalized measure of two cell types co-localization) and G function (Gcross, a measure of two cell types co-localization) were calculated using spatsat^29^ with a radius of 50 μm. Spatial latent Dirichlet allocation (LDA) analysis was done using spatialLDA^30^, using the default radius of 100 μm and 6, 8 or 10 topics for topic modelling. Shorter distances of ∼25 μm may be interpreted as cells nearly or directly touching, while 100 μm represents a distance at which oxygen, nutrients and potentially other molecules diffuse in tissues^31^. Survival analysis was done as described above, using the CyCIF TMA1 dataset as a discovery cohort and the other datasets as the validation cohort. For previously published biomarkers, the validation cohort included patients not used in development of that biomarkers. Specifically, tumor-immune mixing score, which was developed using the MIBI cohort, was validated with the CyCIF and IMC cohorts. Lymphocyte clustering, lymphocyte occupancy and heterotypic neighbor biomarkers were originally developed in external cohorts, so all samples were included in the validation cohort (and no discovery cohort was used). Spatial and survival analysis code, and links to data are available here: https://github.com/engjen/cycIF_TMAs.

## Results

### Analysis of different multiplex imaging datasets produces similar single-cell phenotypes

We generated CyCIF data from two tissue microarrays (TMAs) containing surgical breast cancer specimens and analyzed publicly available breast cancer data from IMC and MIBI platforms generated with similar antigen targets (**Figure 1**). The CyCIF data was comprised of 47 biomarkers imaged at a resolution of 0.325 μm per pixel in entire cores with 1.2 to 1.4 mm diameters, with 1-2 full cores imaged per patient (**Figure 1c top**, S4). The IMC data included 35 biomarkers imaged in the largest square area contained within the TMA cores (from 0.6 - 0.8 mm diameter), at a resolution of 1 μm per pixel (**Figure 1c middle**, S4)^15^. The MIBI data included 36 biomarkers imaged in 0.8 × 0.8 mm square ROIs at a resolution of 0.5 μm per pixel (**Figure 1c bottom**, S4)^13^.

We used the same methods to generate single-cell phenotypes via unsupervised clustering in each platform. In the CyCIF dataset, we generated a 30 k-nearest neighbors (k-NN) graph using a subset of 23 markers common between the CycIF and IMC datasets, plus nuclear area. Markers were scaled by the standard deviation (SD), clipped at 20 SDs, and nearest neighbors were calculated using scanpy^27^. We visualized the k-NN using UMAP embedding, confirming good separation of lineage specific markers, CD31, endothelial, E-cadherin (Ecad), epithelial, Collagen I (ColI), extracellular matrix, vimentin, mesenchymal cells including activated fibroblasts, and CD45, immune infiltrate (S5a). The UMAP visualization showed good mixing of cells from different TMA sources and separation of tumor cells from different breast cancer subtypes (S5b). We used the Leiden algorithm^27^ to cluster the k-NN graph, resulting in 23 cell type clusters (S5c). The mean expression of each biomarker in each cluster was used to annotate cell types (e.g., CD8 T cell, CD4 T cell, luminal ER+ tumor) and lineages (i.e., endothelial, epithelial, fibroblast, immune and other stromal, S5d). The most common cell types included luminal and luminal ER+ tumor, CD4 T cells, vimentin+ fibroblasts and quiescent stroma (S5d). To confirm our cell typing, we performed manual gating on lineage specific markers (S5e). Gating and clustering-based cell lineages localized to similar areas of the UMAP and had 73% accuracy on a single cell level, as calculated using metrics.accuracy_score in scikit-learn^32^ (S5e, f).

In the IMC dataset, we used the same method as in the CyCIF dataset to visualize marker expression, TMA, and subtype and cluster cell types (S6f). We used 21 markers plus nuclear area for clustering, resulting in 24 cell types (S6d). Upon annotation, we found the most common cell types were similar to those in the CyCIF samples, namely luminal, luminal ER+, and ER+ HER2+ tumor, vimentin or fibronectin (FN)+ fibroblasts, quiescent stroma, and T cells (S6d). Clustering and gating-based cell lineages localized to similar areas of the UMAP and had 77% accuracy (S6e, f).

The MIBI panel included more immune-specific markers than the other panels and had 15 markers shared with our CyCIF panel (S4). To generate a dataset in which to audit deeper immune contexture, we clustered on all 33 markers plus nuclear area and eccentricity. Again, we saw good separation of lineage specific markers and Leiden clustering generated 22 cell types of which luminal tumor, fibroblasts, T cells and quiescent stroma were the most common (S7a-c). Gating-based cell types showed good overlap with clustering cell types with 72% accuracy (S7d-e).

### Cell type fractions are similar across platforms

As our most common cell types were similar across platforms, we were encouraged that we could successfully combine the data for cross-platform analysis. We visually validated individual clusters, as well as cell lineages by inspecting the images. In all three platforms, cell lineage identity and spatial distribution matched the underlying imaging data (**Figure 1** c,d).

For further validation, we sought to quantitatively compare the fractions of the five main cell lineages. We calculated the fraction of cells in each lineage for each platform and breast cancer subtype. Both gating and clustering cell types showed high correlation (Pearson R= 0.97 gating and 0.96 clustering) across platforms for ER+ (n=30 CyCIF, 170 IMC) and TNBC (n=59 CyCIF, 50 IMC, 41 MIBI) while HER2+, which had a smaller number of samples (n=8 CyCIF, 22 IMC) had more variability between platforms (**Figure 1** e, f). We did note some platform specific bias; for example, IMC showed a smaller fraction of immune cells defined by clustering in all three subtypes (**Figure 1** e, f). Therefore, when setting high/low cutoffs for cell abundances, we calculated high and low relative to each platform and subtype, as opposed to the whole dataset. Since different antibody clones, probes and imaging systems with were used, resulting in different signal-to-background ratios between platforms, even for the same target, we believe this is a necessary step to account for technical variability.

### Unsupervised clustering defines prognostic tumor subtypes consistent across platforms

For subtyping based on single-cell phenotypes, we calculated the fraction of epithelial or non-epithelial stromal cells in the total epithelial or stromal cells in that sample, respectively. Normalized fractions of epithelial cell types representing greater than 4% of the epithelial compartment were used to cluster ER+ and TNBC patients from all platforms (**Figure 2** a, S8). The resulting seven subtypes included tissues enriched for luminal, basal, luminal ER+, myoepithelial, cytokeratin-low, and proliferating tumor cells, as well as a heterogeneous group not dominated by one phenotype (**Figure 2** a). Each subtype contained a mixture of patients from multiple platforms, with no significant enrichment for any platform (Chi-squared p=0.2 ER+, p=0.52 TNBC, **Figure 2** b, c). The epithelial subtypes present in the ER+ patients were prognostic (log-rank overall-survival (OS) p=0.04, n=162 patients, **Figure 2** d). Cox proportional hazards modelling (CPH) showed that the heterogeneous subtype 6 had a log hazard ratio > 1, indicating poor prognosis (p=0.079, **Figure 2** d). The CPH hazard ratios (HRs) were similarly ordered across the IMC and CyCIF cohorts, with heterogeneous subtype 6 having a HR > 1 in both platforms (S9 a, b). Investigation of the poor prognosis subtype 6 tissues revealed expression of CD44 and EGFR in tumor cells (S9g). Quantification of associated stromal celltypes significant enrichment of CD31+ endothelial cells (Tukey HSD p<0.05) and analysis of epithelial marker expression showed increased CD44 and EGFR expression in subtype 6 (FDR<0.05, **Figure 2** g-h, S9 h). In TNBC patients, the epithelial subtypes were not significantly associated with prognosis (S9c-e).

### Microenvironment subtypes associate with clinical subtype

We then clustered patients based on the stromal cell type fraction in each tissue in a similar manner to the epithelial subtyping above, selecting k=6 for the number of clusters (S10). Since the stromal phenotypes differed across platforms due to different markers, we clustered patients from each platform separately, using the fractions of stromal cell types representing greater than 2% of the stromal compartment. The stromal subtypes were not prognostic, with the exception of the MIBI platform (log-rank=0.003, CPH p=0.056, n=39 patients S10b-g). However, we observed significant correlation between stromal subtypes and clinical subtypes ER+ and TNBC. In the CyCIF cohort, ER+ patients had significantly more of the vimentin+ stromal subtype 0 and less T cell-rich stroma (Chi-squared p=0.098, Bonferroni p-adj<0.05 for subtype 0, n=89 patients, S10e). Similarly, in the IMC cohort, ER+ patients had more Vim+/FN+ fibroblast stromal subtype 0 and significantly less T cell stromal subtype 4 (Chi squared p=0.002, Bonferroni p-adj<0.05 for subtype 4, n=220 patients S10h). Our characterization of stromal subtypes supports the observation that ER+ breast cancer is immune-poor^33^ and shows significant enrichment for vimentin and fibronectin+ fibroblasts relative to TNBC.

### T cells are an independent prognostic factor in TNBC and high proliferation ER+ tumors

Next, we investigated the prognostic value of multiplex imaging-defined cell types within each clinical subtype. We used the CyCIF TMA1 dataset as a training cohort to identify cell types whose fraction of all cells in the tissue were significantly associated with overall survival and cut-offs for high or low abundance. We tested the 0.33, 0.5 and 0.66 quantile to binarize tissues into low and high cell abundance (**Figure 3** a, c, S11a-c). For any cell type showing prognostic significance (log-rank p<0.05), we selected the cut-off with the lowest p-value for validation in the other cohorts (**Figure 3** a, c). Using this methodology, we found that a high abundance of T and B cells were associated with longer OS in TNBC in both the discovery and validation cohorts (validation FDR<0.05, **Figure 3**, a-d).

**Figure 3.**
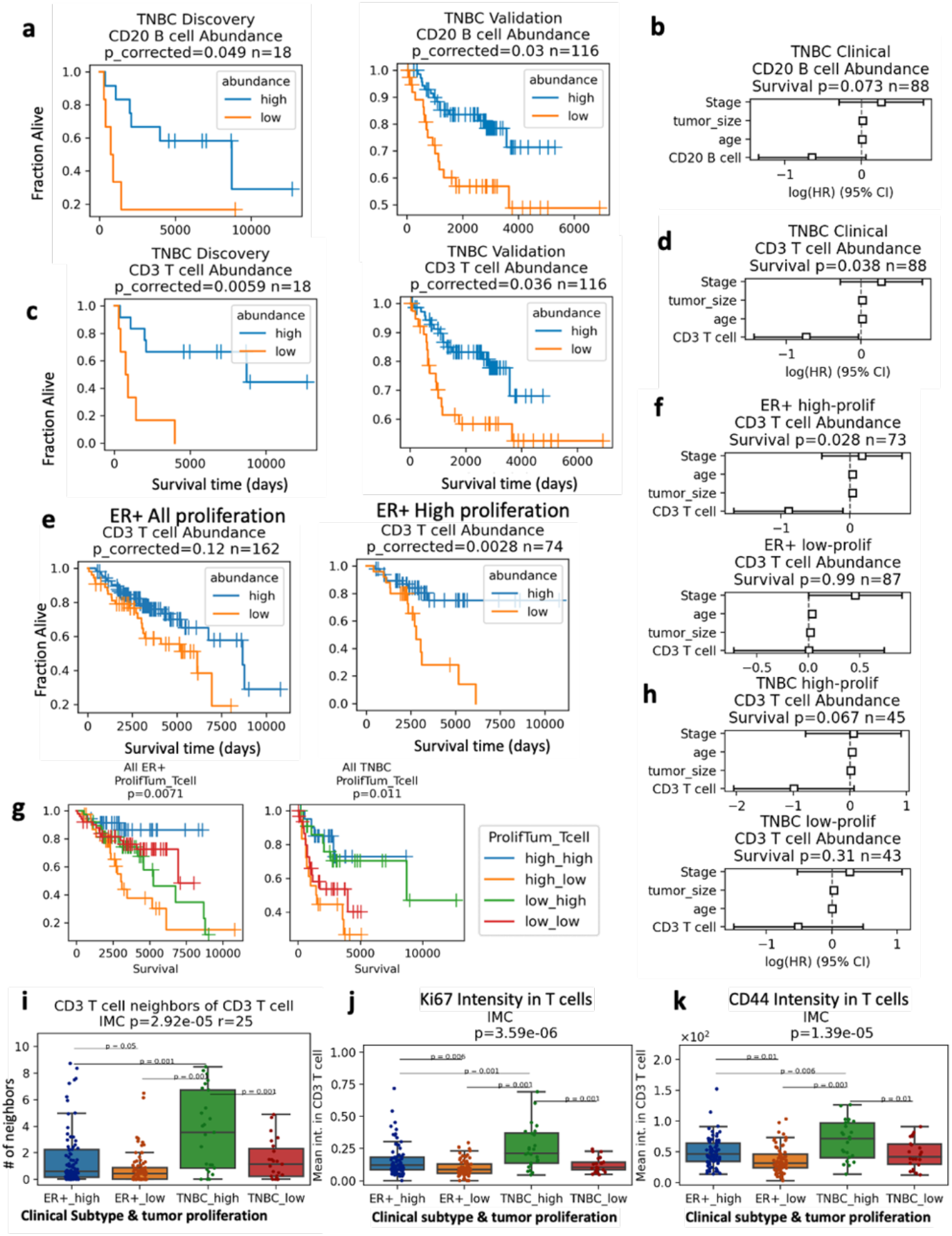
T cell infiltrate has prognostic value in TN and high proliferation ER+ breast cancer. a. Kaplan-Meier analysis of abundance of CD20 B cells versus OS in TNBC discovery (left) and validation cohort (right) b. Multivariate CPH modeling adding patient age, tumor size and stage to CD20 B cell high variable defined in (a). c. Kaplan-Meier analysis of abundance of CD3 T cells versus OS in TNBC discovery (left) and validation cohort (right) d. Multivariate CPH modeling adding patient age, tumor size and stage to CD3 T cell high variable defined in (c). e. Kaplan-Meier analysis of abundance of CD3 T cell versus OS in all ER+ patients (left) and ER+ patients with high (above the median) tumor proliferation (right). f. CPH modeling of CD3 T cell abundance plus clinical variables in high and low proliferation ER+ tumors. g. Kaplan-Meier analysis of all ER+ and TNBC patients stratified into four groups by tumor proliferation and T cell abundance. h. CPH modeling of CD3 T cell abundance plus clinical variables in high and low proliferation TNBC tumors. a-h. All Kaplan-Meier p-values obtained from the log-rank test, validation cohort multiple testing corrected with Benjamini-Hochberg method. CPH modelling p-values for cell type variable given in figure titles. n number of patients given in figure titles. i. Mean number of T cell neighbors (within 25 μm) of T cells in tissues from high and low proliferation ER+ or TNBC tumors in IMC cohort. j. Ki67 intensity indicating proliferation levels of T cells in tissues from high and low proliferation ER+ or TNBC tumors in IMC cohort. k. CD44 intensity in T cells, indicating memory/effector phenotypes in IMC tissues. i-k. Kruskal-Wallis H-test P-value given in figure title. Post-hoc Tukey HSD used for pairwise comparisons between groups.

Biomarkers significant as a single variable were combined with the clinical variables of stage, patient age and tumor size in a multivariate Cox-proportional hazards (CPH) model. In TNBC samples with clinical data, high CD3 T abundance remained significantly associated with longer OS in the multivariate model (CPH p=0.038, n=89, **Figure 3** d). High CD20 B cell abundance was borderline significantly associated with longer OS in the multivariate model (CPH p=0.073, n=89, **Figure 3** b).

CD3 T cells were not associated with OS in the ER+ breast cancer patients (FDR=0.12, n=162), however, in tumors with proliferation above the median, increased CD3 T cells were associated with longer OS (FDR=0.0028, n=74) (**Figure 3** e). Multivariate CPH modelling revealed that high CD3 T cells were independently prognostic for longer OS in high proliferation but not low proliferation ER+ tumors (CPH p=0.028 high proliferation, p=0.99 low proliferation, **Figure 3** f). Separation of tumors by high/low proliferation and high/low T cell abundance showed that in ER+ disease high proliferation, high T cell tumors had the best prognosis, while high proliferation low T cell had the worse prognosis (**Figure 3** g). In TNBC, high- and low-proliferation, high-T cell tumors had similarly good survival, while high- and low-proliferation, low-T cell tumors had similarly poor survival (**Figure 3** f). Multivariate CPH modelling revealed that high CD3 T cells were borderline independently prognostic for longer OS in high proliferation but not low proliferation TNBC tumors (CPH p=0.067 high proliferation, p=0.31 low proliferation, **Figure 3** h). Independent analysis in each platform revealed significant outcome stratification by T cell abundance and proliferation in ER+ patients from the IMC cohort but not the CyCIF cohort (log-rank p=0.028 and 0.4 S12 b), and in TNBC patients from the IMC and CyCIF cohorts but not MIBI cohort (log-rank p=0.056, 0.012 and 0.49 S12 c).

### T cells in patients deriving a survival benefit from infiltration have distinct functional states

In order to identify T cell functional states present in patients deriving a survival benefit from T cell infiltration, we compared T cell spatial localization and marker expression in T cell infiltrated groups with different survival outcomes. There was no significant difference in T cell abundance or T cell to macrophage or endothelial ratio in high versus low proliferation ER+ tumors (S10 e). Interestingly, high-proliferation ER+ tumors in the IMC cohort, which gained survival benefit from CD3 T cells, showed more clustering of CD3 T cells than low-proliferation ER+ tumors, quantified by the number of CD3 T cell neighbors of each CD3 T cell (Tukey HSD p=0.05, **Figure 3** f). High proliferation ER+ tumors in the CyCIF cohort, which did not gain survival benefit from CD3 T cells, did not show increased T cell clustering (Kruskal-Wallis p=0.61, S12 d). Similarly, CD3 T cells in high-proliferation ER+ tumors from IMC cohort had higher levels of the proliferation marker Ki67 and the memory/effector marker CD44 than in low proliferation ER+ tumors, indicating a more activated functional state (Tukey HSD p=0.006 and 0.01, **Figure 3** i-k). In the CyCIF cohort, similar differences in Ki67 and CD44 expression were observed between ER+ and TNBC subtypes, consistent with an activated T cell state correlating with a survival benefit derived from increased T cell infiltration (S12 f). High-proliferation TNBC, in which T cells independently predicted overall survival in the multivariate CPH model (**Figure 3** j, k, also showed increased levels of PD-1, FoxP3, IDO and Lag3 expression in T cells, consistent with upregulation of negative feedback checkpoints following immune activation (S12 f-g). Epithelial cells in high-proliferation TNBC had increased expression of the antigen presentation molecule HLA-Class-1 and immune checkpoint PD-L1 (S10 h).

### Analysis of tumor-immune interactions reveals conserved spatial biomarkers

Intrigued by the finding that T-cells in high-proliferation tumors had increased T cell neighbors and were associated with a survival benefit (**Figure 3** i), we leveraged our discovery and validation cohort analysis to systematically investigate spatial tumor-immune interactions as biomarkers in breast cancer. We calculated the number of immune and tumor neighbors within proximity of each other to derived previously published biomarkers including mean neighbor counts^12,14^, tumor-immune mixing score^13^, lymphocyte clustering and lymphocyte occupancy^16^. We also used common statistical methods for quantification of spatial correlation (Ripley’s L, Kcross and Gcross functions^29^). Analysis of cell lineage proximity revealed that stromal (non-fibroblast, non-immune, non-endothelial) neighbors of epithelial cells predicted longer RFS in the discovery (log-rank p=0.018) and validation ER+ cohorts (log-rank FDR=0.028), and independently predicted RFS in a multivariate CPH model (CPH p=0.02, **Figure 4** a-c). Immune neighbors of immune were associated with longer OS in both TNBC cohorts (validation log-rank FDR=0.019), but not in the multivariable model (CPH p=0.14, **Figure 4** c-d). Analysis of cell type neighbors showed increased macrophage neighbors of tumor and increased B cell neighbors of T cells predicted longer RFS in TNBC in both cohorts (validation FDR=0.05) and remained significant the multivariate model (CPH p=0.047 Macrophage-tumor, p=0.017 B cell-T cell, **Figure 4** e-h). Additional spatial metrics, including Ripley’s L, Kcross and Gcross, did not yield any significant validated biomarkers (S13a, g).

**Figure 4.**
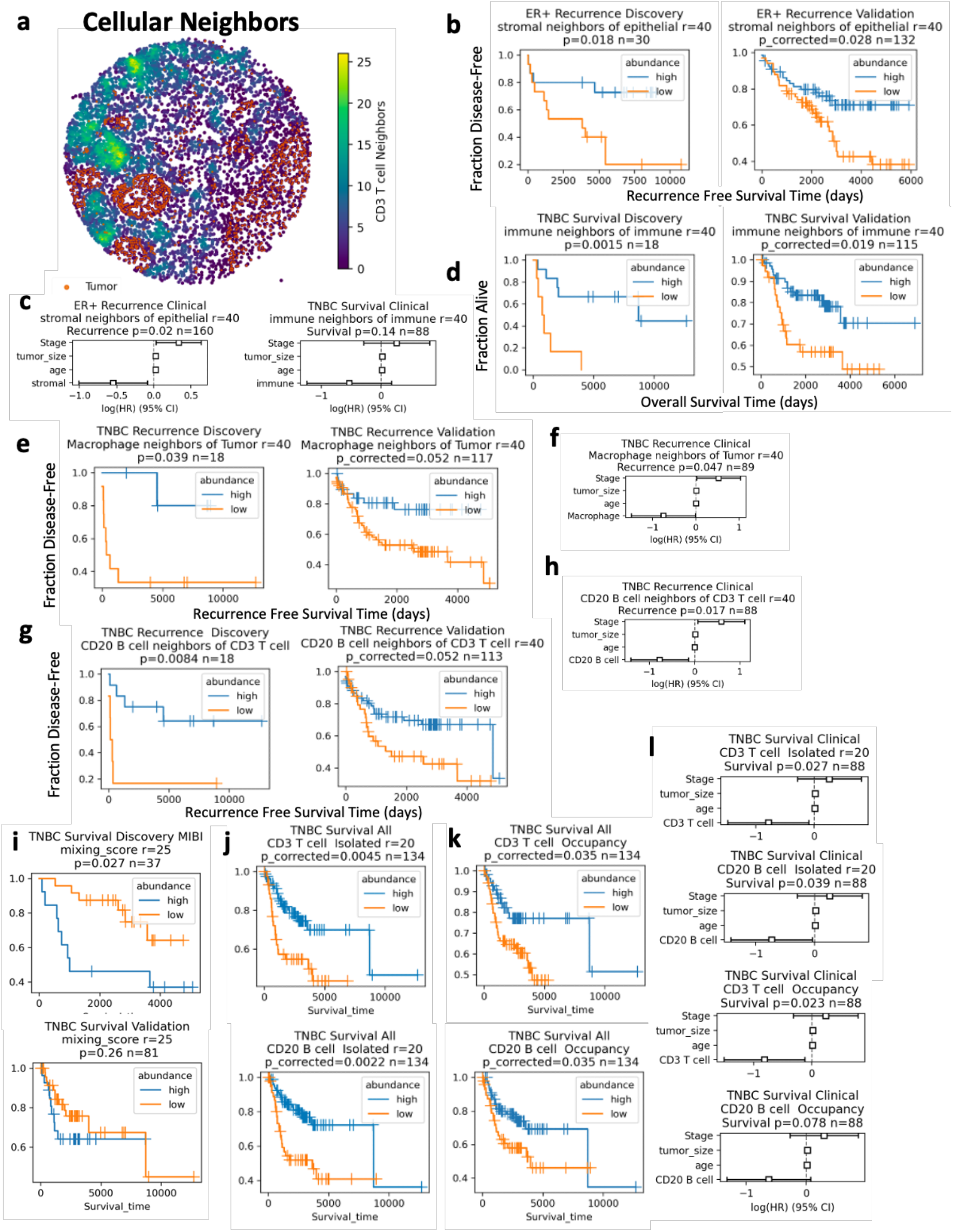
Prognostic tumor-immune spatial correlations in breast cancer cohorts. a. Spatial locations of cells in a TMA core with each cell colored by number of CD3 T cell neighbors in a 40 μm radius. Tumor cells shown in orange. b. Recurrence free survival (RFS) Kaplan-Meier (K-M) analysis of ER+ patients with high versus low stromal neighbors of epithelial cells in a 40 μm radius in the discovery (left) and validation (right) cohorts. c. Left - Multivariate CPH modeling of ER+ patient RFS adding patient age, tumor size and stage to the stromal neighbors of epithelial variable defined in (b). Right - multivariate CPH modeling of TNBC patient OS versus immune neighbors of immune cells. d. Overall survival (OS) K-M analysis of TNBC patients with high versus low immune neighbors of immune cells in a 40 μm radius in the discovery (left) and validation (right) cohorts. e & g. RFS K-M analysis of TNBC patients with high versus low macrophage neighbors of tumor (e) or B cell neighbors of T cells (g) in a 40 μm radius in the discovery (left) and validation (right) cohorts. f & h. Multivariate CPH modeling of TNBC patient RFS versus macrophage neighbors of tumor (f) or CD20 B cell neighbors of CD3 T cells (h). i. K-M analysis of TNBC OS stratified by tumor immune mixing score developed by Keren et al. in the MIBI cohort (top) and validation cohort (i.e. CyCIF and IMC; bottom). j. K-M analysis of TNBC OS stratified by isolated T (top) or B lymphocyte (bottom) metric developed by Wortman et al. k. K-M analysis of TNBC OS stratified by T (top) or B lymphocyte (bottom) occupancy metric developed by Wortman et al. l. CPH modeling of Wortman et al. metrics from j and k. a-l. All Kaplan-Meier p-values obtained from the log-rank test. CPH modelling p-values for spatial variable given in figure titles. N number of patients given in figure titles..

Previously, Ali *et al*.^14^ showed that heterotypic neighborhoods of myofibroblasts, fibroblasts, cytokeratin low tumor cells, and vimentin+ Slug-macrophages were associated with poor outcome and homotypic neighbors of fibroblasts and myofibroblasts were associate with good outcome in all breast cancer subtypes. We tested the prognostic value of heterotypic and homotypic neighbors of fibroblast subsets, CK low tumor and macrophages but did not find the significant association with survival (log-rank FDR>0.3, S13 b, c). Previously, Keren *et al*.^13^ showed that a high tumor-immune mixing score was associated with poor survival in TNBC. Encouragingly, we were able to reproduce the prognostic value of the mixing score in the MIBI cohort, where it was developed (log-rank p=.027, **Figure 4** i). However, in a validation cohort containing CyCIF and IMC patients, the mixing score was not prognostic (log-rank p=.26, **Figure 4** i), nor was it independently prognostic in samples with clinical outcome (CPH p=0.4, S13 e). Previously, Wortman *et al*.^16^ showed that TNBC with longer RFS had more spatially dispersed lymphocytes, as defined by isolated lymphocytes at <5 per 20 um radius, and an increased tissue area occupied by lymphocytes. Our analysis supported the findings of Wortman et al., with greater numbers of isolated T cand B cells and higher T and B cell area occupancy in tissue associated with longer RFS (log-rank FDR <0.05, **Figure 4** j, k). Isolated lymphocytes and lymphocyte occupancy also remained prognostic in the multivariable model in TNBC (CPH p<0.05, except CD20 B cell occupancy p=0.078, **Figure 4** l).

To identify spatial metrics which provided additional information beyond abundance, we calculated the Pearson correlation between all our spatial metrics and cell type abundance within each patient’s tissue (S14). Stromal neighbors of epithelial (good prognosis in ER+) correlated with quiescent stroma abundance and macrophage neighbors of tumor (good prognosis in TNBC) correlated with macrophage abundance (S14). CD20 B cell neighbors of T cells, isolated lymphocytes and lymphocyte occupancy (good prognosis in TNBC) correlated with each other and T and B cell abundance (S14). The tumor-immune mixing score was positively correlated with tumor abundance and negatively correlated with immune abundance (S14). Many of the Kcross and Ripley’s L function results correlated with each other and were not as strongly correlated with abundance (S14).

### Neighborhood analysis reveals multicellular spatial biomarkers

Finally, we analyzed multicellular spatial neighborhoods by considering all stromal cells within 100 μm radius of each tumor cell. We used spatial latent Dirichlet allocation (LDA) to model the neighborhood around each tumor cell as a combination of topics, utilizing a spatial parameter to increase the likelihood that adjacent cells share the same topics^30^. In LDA analysis, each topic can contain multiple cell types and each cell type can be present in multiple topics. In our CyCIF data, for example, topic-0 in TNBC tissues was enriched in macrophages, vimentin+ fibroblasts and CD4 T cells, while CD4 T cells are found in topic-0, 4, 5 and 6 (**Figure 5** a). After topic modelling, K-means clustering was run on the single-cell topic matrix to define “tumor neighborhood” clusters which contained one or more topics (**Figure 5** b). Clustering the topic matrix rather than the neighbor count matrix is believed to be less sensitive to noise^34^ and allows for smooth transitions between neighborhoods as opposed to arbitrary cutoffs^30^. We did observe transitioning/mixed neighborhoods within both TNBC and ER+ cluster results (**Figure 6** b, d, S15). The spatial LDA neighborhood clusters were annotated based on their topics and examination of the images showed neighborhoods reflected the spatial distribution of the markers in the tissue (S15a-d). A TNBC tissue in our CyCIF cohort, for example, showed tumor cell neighborhoods with more T cells (blue) on the tumor margin, with adjacent macrophage-rich tumor neighborhoods (purple) (S15 a-b). These neighborhoods transition into a mixed neighborhood (brown), and finally a vimentin+ FB neighborhood (green) distant from the infiltrating T cells (S15 a-b). An ER+ tissue showed quiescent stroma tumor cell neighborhoods in the tumor core, vimentin+ FB neighborhoods on the tumor margin and T cell neighborhoods in tumor nests isolated in the stroma (S15 c-d). Similar neighborhoods were identified in the IMC TNBC and ER+ cohorts (S15 e-h) but the MIBI cohort was excluded from this analysis due to lower stromal cell type overlap (**Figure 1**). For survival analysis, we used the CyCIF TMA1 as a discovery cohort and CyCIF TMA2 plus IMC patients as a validation cohort. Due to fewer numbers of patients, we used an alpha of 0.1 rather than 0.05 to report findings. Increased Vimentin+ FB neighborhoods around tumor cells were associated with shorter OS in both TNBC cohorts (validation log-rank p=0.07, **Figure 5** e). Increased vimentin+ fibroblast neighborhoods were associated with shorter OS and RFS in the multivariate model (CPH p=0.049 OS, 0.053 RFS **Figure 7** f, S14b). Interestingly, vimentin+ fibroblast abundance alone was not prognostic, although neighborhood abundance was correlated with fibroblast abundance (S16). In ER+ tumors, increased mixed fibroblast neighborhoods containing vimentin-positive and negative fibroblasts around tumor cells were associated with shorter OS (validation log-rank p=0.088), and trended significant in the multivariable model (CPH p=0.087 OS, p=0.046 RFS, **Figure 5** g-h). Finally, similar to other groups^34^, we found that directly clustering the neighborhood counts using Kmeans (rather than running LDA and clustering the topics) did not result in robust prediction of prognosis (S17).

**Figure 5.**
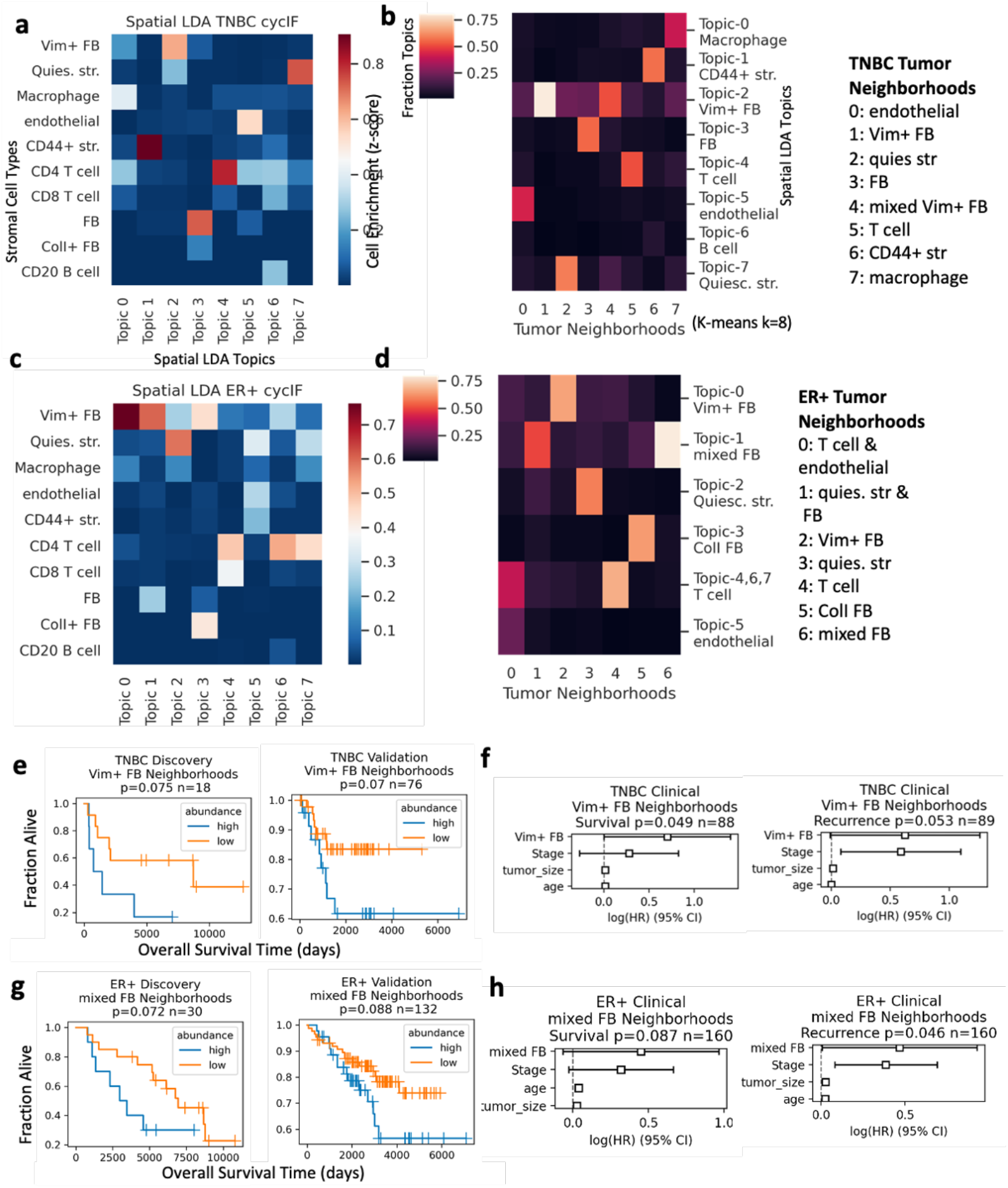
Prognostic multicellular neighborhoods surrounding tumor cells modeled with spatial latent Dirichlet allocation. a. Heatmap of stromal cell enrichment in spatial latent Dirichlet allocation (LDA) topic models of 100 μm TNBC tumor neighborhoods in the CyCIF cohort. b. Heatmap of fraction of each topic in each neighborhood cluster resulting from K-means clustering (k=8) of spatial LDA topics from (a). c-d. heatmaps as defined in a and b, for ER+ tumors from the CyCIF cohort.e. Kaplan-Meier (K-M) analysis of overall survival (OS) versus high or low vimentin+ fibroblast neighborhoods in TNBC tissues in discovery (left) and validation cohorts (right). p-values obtained from the log-rank test; n given in figure titles. f. CPH modelling of OS and recurrence-free survival (RFS) with clinical variables plus spatial LDA neighborhoods from (e). p-values and n number of patients given in figure title. g. K-M analysis of OS versus high and low mixed fibroblast neighborhoods in ER+ tissues in the discovery (left) and validation cohorts (right). h. CPH modeling of OS and RFS for mixed fibroblast neighborhoods in ER+ tumors.

### Tumor phenotypes correlate with stromal cell abundance and spatial neighborhoods

We hypothesized that there would be significant correlation between tumor cell types and the surrounding stromal cell neighborhoods, correlations that could shed light on biologically and clinically relevant tumor-stroma crosstalk. First, we visualized a matrix of pairwise correlation between epithelial and stromal cell fractions and spatial LDA neighborhoods across subtypes (S16a, c). Epithelial cell types were inversely correlated with each other, indicating most tumors had just one main epithelial cell type (S16a, c). The exception was luminal tumor, which correlated with cytokeratin low tumor in ER+ breast cancer, indicating mixing of these phenotypes within the same tissues (S16c).

Immune cells exhibited distinct tissue-level correlations in the different subtypes. In TNBC, T cells correlated with proliferating tumor, and macrophages correlated with CD4 T cells (S16a). In ER+ breast cancer, T cells correlated with B cells while proliferating tumor, macrophages and endothelial cells were correlated with each other but not T cells (S16c). Vimentin+, FN+, and ColI+ fibroblasts, as well as quiescent stroma were inversely correlated with immune cells (S16a, c). In both subtypes, spatial LDA neighborhoods correlated strongly with the abundance of their respective stromal cell types; however, neighborhoods showed unique correlations to other cell types present. For example, proliferating and luminal ER+ tumor cell abundances did not correlate with total T cell abundance in ER+ breast cancer, but they did correlate with the fraction of T cell neighborhoods (S16c). In TNBC, vimentin+ and fibronectin+ fibroblast abundances were not correlated, but their respective neighborhoods were inversely correlated, suggesting tissue exclusivity for a single fibroblast phenotype near tumor cells (S16a, b). Therefore, although spatial neighborhoods tend to correlate with cell abundance, they do reveal unique features of tumor-stromal organization in tissues.

## Discussion

Our approach of combined analysis across multiple platforms shows the power of our methods for biomarker discovery. We were able to incorporate analysis of two publicly available imaging datasets with our own CyCIF data for efficient discovery of robust biomarkers.

We utilized our validated method for CyCIF staining and image processing^18^ to generate multiplex imaging data of 42 markers in a single tissue section from two TMAs with clinical follow-up. Our dataset alone represents a valuable new clinical cohort that provides improved plex, resolution, and ROI size compared to previously published datasets^13,15^. We then developed an analysis pipeline (https://github.com/engjen/cycIF_TMAs) to generate single cell phenotyping data from our CyCIF dataset and two publicly available datasets^13,15^. The advantage of using our pipeline for image processing is the development of smoothing algorithms so that pixelated image data can be segmented with deep-learning models trained on higher resolution images and an algorithm to match nuclear and cell segmentation results from separate deep-learning segmentation models to extract features from subcellular compartments such as the nucleus and cytoplasm (S1b). Using our methods, we generated single cell data that produced a high correlation between cell type fractions across cohorts from the same breast cancer subtype and different platforms (**Figure 1**).

Additionally, we were able identify similar epithelial phenotypes across platforms and cluster patient data from all platforms to separate seven epithelial subtypes that did not have a platform-specific bias (**Figure 2**). Our subtypes were similar to the intrinsic breast cancer subtypes^35^, including a luminal ER+ luminal A-like group with good prognosis and a cytokeratin low group previously shown to share features with luminal B tumors^14^. Triple-negative breast cancers also fell into categories similar to those defined by gene expression profiling^36^, including a highly proliferative, basal-like 1 (BL1)-like group, a luminal androgen receptor (LAR)-like group with luminal epithelial phenotypes, and a group with a basal/myoepithelial phenotype reminiscent of the basal-like 2 (BL2) group. We also identified a heterogeneous subtype with low cytokeratin and elevated CD44 expression that may represent tumors with mesenchymal features. Jackson *et al*., identified a similar single cell pathology cluster of hormone-receptor positive mixed tumors with poor prognosis^15^. In our analysis, the ER+ tumors in the heterogenous subtype had poor prognosis and showed increased angiogenesis. An EMT-program in breast cancer cells is linked to increased vascular endothelial growth factor A expression, increasing angiogenesis and the capacity for tumor initiation^37^, a mechanism that could explain these correlated tumor and stromal phenotypes and their association with poor outcome.

Tumor infiltrating lymphocytes have been linked to good prognosis in TNBC^38^, and we confirmed that T and B cells are independently prognostic in TNBC in the multiplex imaging datasets analyzed herein. Previous gene expression profiling studies link productive anti-tumor immunity and tumor proliferation. Nagalla *et al*. found that immune signatures were prognostic solely in breast cancer patients with the highest proliferation gene expression^39^. Subsequently, the same group showed that immune gene signatures were prognostic in highly proliferative basal-like, HER2-enriched and luminal B subtypes, but not those with low proliferation^40^. Similar to this previous work, we showed that CD3 T cells were independently prognostic specifically in high-proliferation ER+ and TNBC tumors.

Our analysis of immune functional states showed increased T cell proliferation, activation and checkpoint molecules and epithelial antigen presentation in high-proliferation tumors, consistent with IFNγ pathway activation. Consistent with our analysis, gene network analysis previously showed activation of TNFα/IFNγ signaling pathways in tumors with productive anti-tumor immunity and TGF-β, an immunosuppressive cytokine, in tumors with unproductive anti-tumor immunity^40^. TGF-β also has anti-proliferative effects and is associated with good outcome in ER+ breast cancer cohorts^41^, suggesting that it could mechanistically link lower proliferation rates with immunosuppression and represent a rational drug combination with immune checkpoint targeting^42^.

More recently, the Thomas *et al*. showed that immune gene signatures were prognostic exclusively in tumor-mutation burden (TMB)-high breast cancer tumors^43^. Thirty-seven percent of basal-like tumors had high TMB, while only 11.5% of luminal A tumors did^43^, explaining the poor immunogenicity of the latter subtype. Together, these data point to a model of high TMB correlating with high-proliferation and both linked to productive anti-tumor immunity. It had been hypothesized that oncogenes driving sustained proliferation also induce DNA replication stress, which generates genomic instability and would increase TMB^44^. In summary, TMB provides a mechanistic link between proliferation and anti-tumor immunity and should be investigated in future studies. Furthermore, our analysis shows enrichment of potential immune checkpoint targets in high-proliferation breast cancer, including PD1, Lag3, IDO and PD-L1 elevation (S12).

One of the main goals of this study was to provide methods and a framework for robust identification of spatial biomarkers. Our use of external cohorts to validate biomarkers discovered in our CyCIF data increases our confidence in biomarker identification. In ER+ breast cancer, we found that increased stromal neighbors of tumor correlated with better prognosis, similar to previous studies showing survival benefit for high stroma in ER+ tumors^45^. We found that macrophage proximity to tumor was associated with good prognosis in TNBC, which is surprising given previous publications. Specifically, tumor associated macrophages (TAMs)^46^ were associated with shorter overall survival in a cohort of ER+ and ER-patients, but the prognostic value of macrophages specifically in TNBC was not investigated. Furthermore, Medrek *et al*.^47^ found that CD68+ macrophages in close proximity to tumor cells were not associated with poor survival, but those out in the stroma were, suggesting that it may be difficult to compare our metric of macrophage-tumor neighbors in a 40 μm radius with previous studies and further investigation is warranted. Finally, numerous immune-related spatial biomarkers, including immune-immune proximity, B cell-T cell proximity, isolated lymphocyte abundance and lymphocyte occupancy were associated with good prognosis in TNBC, supporting a model of productive anti-tumor immunity in the triple-negative subtype. Encouragingly, our results for the prognostic value of lymphocyte spatial metrics were similar to those Wortman *et al*.^16^ discovered in a different TN cohort.

We utilized spatial LDA modelling to analyze multicellular neighborhoods of stromal cells surrounding tumor cells. We identified a neighborhood enriched for vimentin+ fibroblasts that was independently associated with shorter survival in TNBC. Given the high levels of vimentin and low levels of alpha-SMA, these cells may have an inflammatory phenotype similar to CAFs that differentiate under TNFα + IL-1β stimulation^48^. Interestingly, TNFα + IL-1β have been shown to stimulate pro-metastatic chemokine expression (CXCL8, CCL2 and CCL5) and aggressive characteristics in TNBC cell lines, mediated in part by direct CAF-tumor cell contact in co-cultures^49^, consistent with close proximity between poor-prognosis CAFs and tumor cells in spatial LDA neighborhoods.

The limitations of our study include different antibody probes and imaging systems resulting in different signal-to-background ratios for biomarkers across platforms. Therefore, we relied on matching hand-annotated clusters across platforms for our integrated analysis. Since our annotations may not correspond to the same cell types in each platform, this introduces uncertainty. Some well-defined phenotypes, such as T cells and proliferating tumor, are relatively straightforward. However, variable performance of antibodies, such as anti-ER, for example, could lead to variability in the classification of phenotypes such as luminal ER+ versus luminal tumor across platforms. To correct for platform-specific bias in cell types, we binarized patients into high/low expression within each subtype and platform for survival analysis. However, such binarization may not reflect underlying heterogeneity in quantitative biomarker abundance.

Another limitation of our study is the use of 1-2 TMA cores per patient for analysis. It has been shown that a limited number of TMA cores (≤3) are needed to binarize patients in to high and low TILs, although a larger number of cores (≥11) are needed to accurately estimate the mean TIL abundance^50^. Our survival analysis relied on binarizing patients; therefore, the use of TMAs may be appropriate in this context. Lacking large tissue sections, we had no way to quantify sensitivity of spatial metrics to sampling bias introduced by analyzing small ROIs, but our cross-cohort analysis revealed a lack of generalization of spatial biomarkers in several cases (S13a). Performance of spatial biomarkers could potentially be improved by sampling larger tissue areas, although increased heterogeneity in large sections could introduce noise. The optimal balance between the area analyzed in each tissue and number of patients included for estimation of prognostic tumor microenvironment composition and spatial architecture remains an open and important question in the field.

Overall, our spatial analysis supports the utility of spatial information in uncovering novel biomarkers of patient outcome in breast cancer. The tools developed in this study can be utilized to analyze additional cohorts for further characterization of biomarkers in breast cancer and other tumor types.

## Acknowledgements

We would like to thank Dr. Joe Gray for invaluable advice on analysis and feedback on the manuscript. We would also like to thank Drs. Sandra Rugonyi, Sara Courtneidge, Young Hwan Chang, and Pepper Schedin for feedback on earlier versions of this work. We would like to thank Damir Sudar for technical support related to image storage and Elliot Gray for sharing his neighbor counter script. We sincerely appreciate Drs. Clare Yu, Peter Lee and Julie Wortman for sharing their lymphocyte occupancy code. We appreciate the sample scanning assistance of both Drs. Stefanie Kaech Petrie and Crystal Chaw at the OHSU Advanced Light Microscopy Core. This work was supported by funding from NIH/NCI U54 CA209988, the Prospect Creek Foundation, and OHSU Foundation. We received the services from the Knight Cancer institute Histopathology Shared Resource and Advanced Light Microscopy Core, supported by the Cancer Center Support Grant (NIH/NCI P30CA69533). Additionally, J.A.P acknowledges support from NIH P50 CA098131 and Komen SAB210301.

## Author Contributions

J.E., S.L.G., and K.C. conceived of the project. M.S., B.C., P.G., and J.A.P. collected patient tissues, clinical outcome data, and constructed the TMAs. Z.H. performed staining and imaging experiments. J.E. and E.B. did single-cell image processing. J.E. ran the analysis, drafted the manuscript, and prepared the figures. S.L.G., K.C., E.B., B.C. and J.A.P. edited the manuscript.

