## Supplemental Figures for "Robust biomarker discovery through multiplatform multiplex image analysis of breast cancer clinical cohorts"

#### LIST OF SUPPLEMENTARY FIGURES

Supplementary Figure S1. De-noising and Segmentation Optimization.
Supplementary Figure S2. Mesmer versus Watershed Segmentation.
Supplementary Figure S3. ER Quality Control.
Supplementary Figure S4. Comparison of breast cancer MXI panels and dataset sizes.
Supplementary Figure S5. Single Cell Analysis of CyCIF data.
Supplementary Figure S6. Single Cell Analysis of IMC data.
Supplementary Figure S7. Single Cell Analysis of MIBI data.
Supplementary Figure S8. Normalization of epithelial fractions across platforms.
Supplementary Figure S9. Prognostic Value of Epithelial Subtypes.
Supplementary Figure S10. Prognostic Value and Clinical Subtype Correlation of Stromal Subtypes.
Supplementary Figure S11. Single Variable Prognosis in separate cohorts.
Supplementary Figure S12. Prognostic value of proliferation and T cell abundance.
Supplementary Figure S13. Prognostic value of tumor immune spatial metrics.
Supplementary Figure S14. Correlation of Spatial Metrics and tissue composition.
Supplementary Figure S15. Spatial LDA Tumor Neighborhoods.
Supplementary Figure S16. Correlation of tumor neighborhoods and tissue composition.
Supplementary Figure S17. Neighborhoods defined by directly clustering the cell counts.

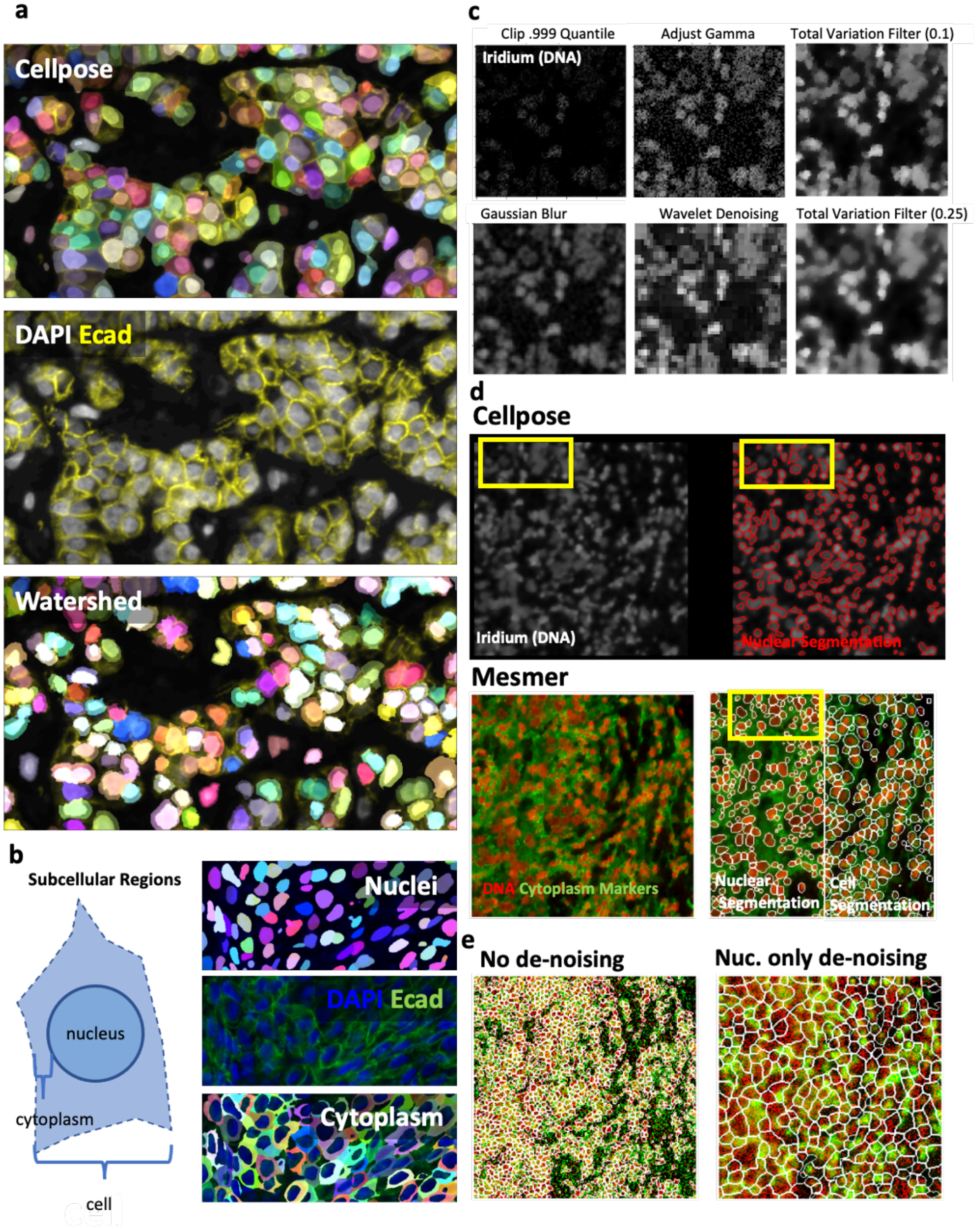


#### S1. De-noising and Segmentation Optimization.

a. CyCIF Cellpose nuclear and cell segmentation (top) versus watershed segmentation (bottom). b. Construction of cytoplasm mask from cell and nuclear masks after matching with mplexable. c. Image processing steps tested for the IMC denoising pipeline. d. IMC Cellpose segmentation (top) versus Mesmer segmentation (bottom). e. IMC Mesmer segmentation of gamma-adjusted only image of ROI shown in (d) (left), and of de-noised nuclear channel with no cytoplasm denoising (right).

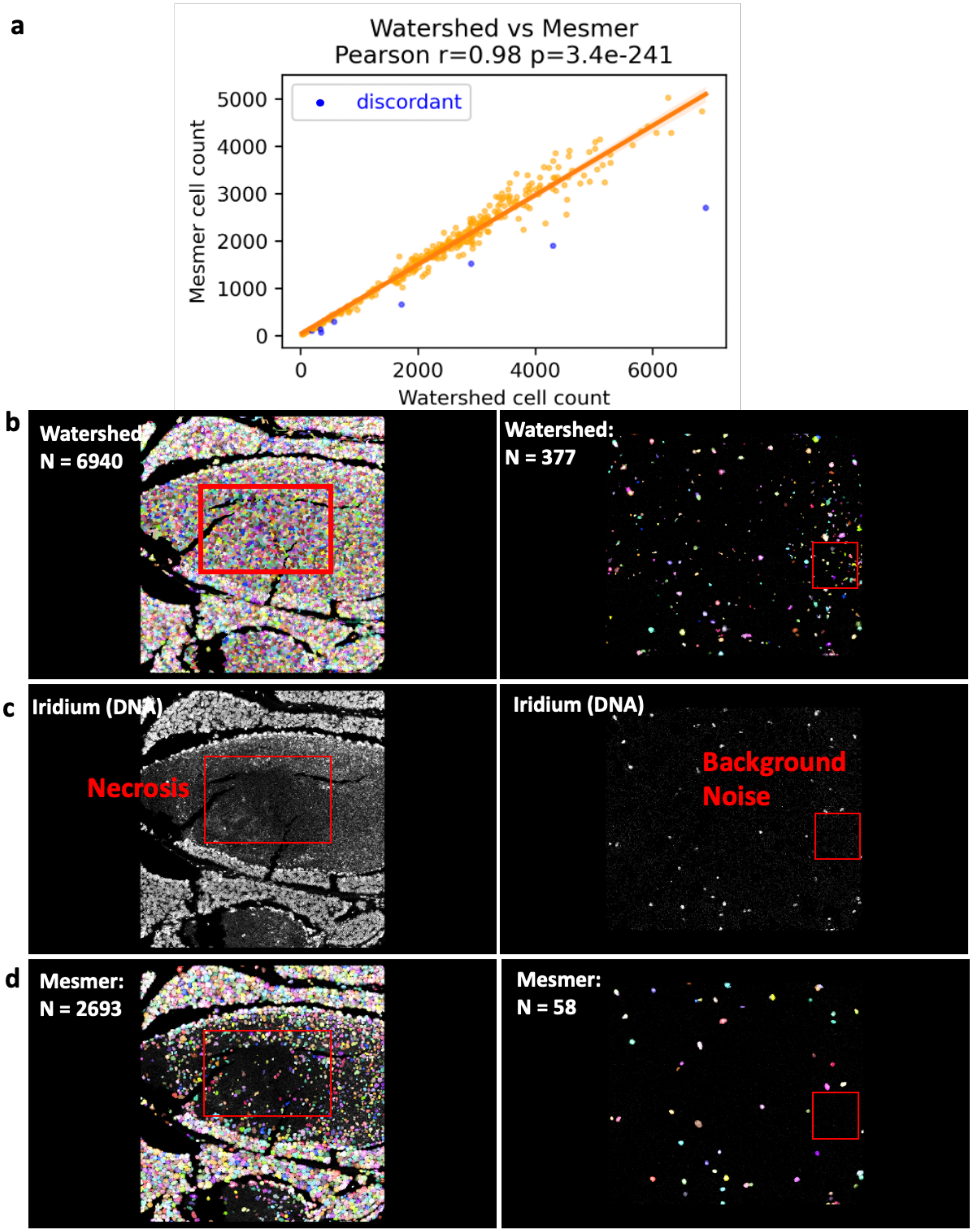


**S2. Mesmer versus Watershed Segmentation.**

a. a. Pearson correlation between cell counts of watershed versus Mesmer segmentation in IMC tissues. Tissues with >50% change shown in blue. b. Watershed segmentation of two selected tissues with discordant cell counts between segmentation methods. Red boxes indicate areas of over segmentation. c. DNA channel of selected tissues. d. Mesmer segmentation of selected tissues.

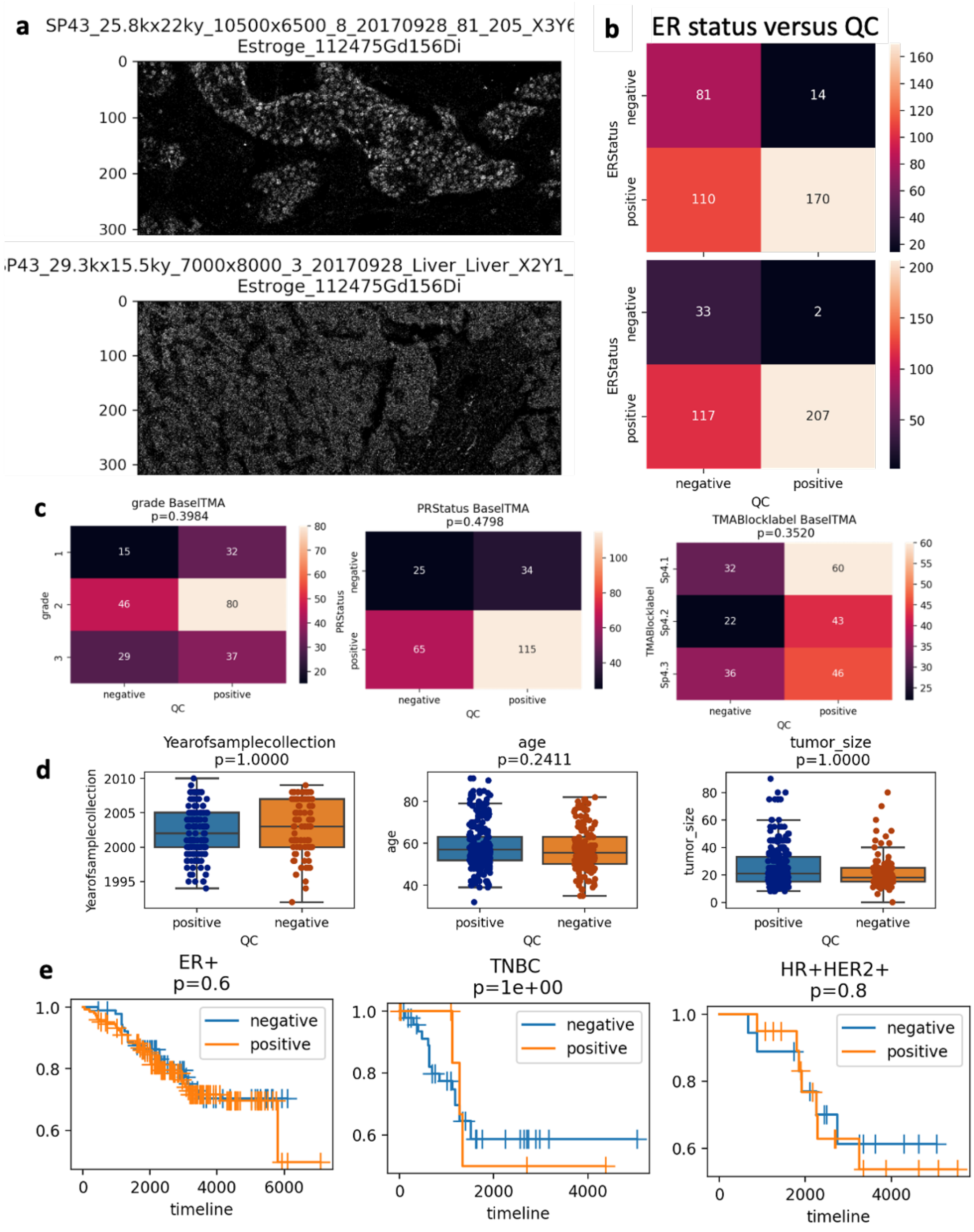


### S3. ER Quality Control.

a. Representative IMC images of estrogen receptor channel QC images sorted for positive (top) and negative (bottom) nuclear ER staining. b. Image ROIs classified by annotated ER status (y-axis) versus our QC call for ER positive or negative (x-axis) in two IMC TMAs. c. Grade, PR status and TMA block versus QC calls, p-values from Chi-squared analysis shown in figure title. b-c. N number of ROIs in each category annotated on heatmap cells. d. Year of sample collection, patient age and tumor size versus QC status of each ROI, p-values for Wilcoxon-rank sum test shown in figure title. e. Kaplan-Meier curves for OS versus QC status, p-values from log-rank test.

a

| CycIF Antigen | Description | Antigen | Description |
|---|---|---|---|
| Cytokeratins | | Immune Context | |
| CK5 | Basal CK | CD45 | pan-Immune |
| CK7 | Luminal CK | CD3 | T Cells |
| CK8 | Luminal CK | CD20 | B Cells |
| CK14 | Basal CK | CD8 | Cytotoxic T cell |
| CK17 | Basal CK | CD4 | Helper T cell |
| CK19 | Luminal CK | CD68 | Macrophage |
| Adhesion Molecules | | PD1 | T cell exhaustion |
| Ecadherin | Cell Adhesion | FoxP3 | Regulatory T cell |
| CD44 | Cell Adhesion | GRNZB | Cytotoxicity |
| Hormone Receptors | | Hypoxia and Metabolism | |
| ER | Estrogen receptor | HIF1a | Hypoxia |
| PR | Progesterone Receptor | Glut1 | Glucose Transport |
| AR | Androgen Receptor | CoxIV | Mitochndria |
| RTK Signaling | | Cell Growth and Division | |
| HER2 | RTK | Ki67 | Proliferation |
| EGFR | RTK | PCNA | Proliferation |
| Endothelial | | p-HH3 | Mitosis |
| CD31 | Endothelial | p-S6 | Growth |
| PDPN | Lymphatic Endothelial | p-RB | Cell Division |
| Stromal Markers | | Nuclear | |
| SMA | Myoepithelial cells | Lamin AC | Nuclear Membrane |
| Vimentin | Mesenchymal Cells | Lamin B1 | Nuclear Membrane |
| Collagen IV | Basement Membrane | Lamin B2 | Nuclear Membrane |
| Collagen I | Extracellular Matrix | Cell Death | |
| Epigenetic Mark | | cleaved PARP | Apoptosis |
| H3K27 | Gene Repression | | |
| H3K4 | Gene Accessibility | | |

| MIBI Antigen | Description | Antigen | Description |
|---|---|---|---|
| Cytokeratins | | Immune Context | |
| CK6 | Basal CK | CD45 | pan-Immune |
| CK17 | Basal CK | CD3 | T Cells |
| pan-CK | Luminal & Basal | CD20 | B Cells |
| Adhesion Molecules | | CD8 | Cytotoxic T cell |
| Beta catenin | Cell Adhesion | CD4 | Helper T cell |
| Transcription Factors | | CD68 | Macrophage |
| p53 | Tumor suppressor | PD1 | Immune regulation |
| RTK Signaling | | FoxP3 | Regulatory T cell |
| EGFR | RTK | CD45RO | Memory T cells |
| Endothelial | | CD56 | NK cells |
| CD31 | Endothelial | HLA Class 1 | Antigen Presentation |
| Stromal Markers | | HLA DR | Antigen Presentation |
| SMA | Myoepithelial cells | CD11c | Dendritic cells |
| Vimentin | Mesencymal Cells | CD11b | pan-Myeloid |
| Epigenetic Mark | | CD138 | Plasma cells |
| H3K27me3 | Gene Repression | CD16 | NK cells |
| H3K9ac | Gene Accessibility | CD209 | Antigen Presentation |
| Cell Growth and Division | | IDO | Immune regulation |
| Ki67 | Proliferation | Lag3 | Immune regulation |
| p-S6 | Growth | MPO | Neutrophils |
| | | PDL1 | Immune regulation |
| | | CD63 | Monocytes |

| IMC Antigen | Description | Antigen | Description |
|---|---|---|---|
| Cytokeratins | | Immune Context | |
| CK5 | Basal CK | CD45 | pan-Immune |
| CK7 | Luminal CK | CD3 | T Cells |
| CK8/18 | Luminal CK | CD20 | B Cells |
| CK14 | Basal CK | CD68 | Macrophage |
| CK19 | Luminal CK | Transcription Factors | |
| AE1/AE3 | Pan-CK | p53 | Tumor Suppressor |
| Adhesion Molecules | | cMyc | Proto-Oncogene |
| E/P-cadherin | Cell Adhesion | GATA3 | Luminal TF |
| CD44 | Cell Adhesion | Twist | EMT TF |
| Hormone Receptors | | Slug | EMT TF |
| ER | Estrogen receptor | Hypoxia and Metabolism | |
| PR | Progesterone Receptor | HIF1a | Hypoxia |
| RTK Signaling | | Glut1 | Glucose Transport |
| HER2 | RTK | CoxIV | Mitochondria |
| HER2 | RTK | Cell Growth and Division | |
| Endothelial | | Ki67 | Proliferation |
| vWF | Endothelial | p-HH3 | Mitosis |
| CD31 | Endothelial | p-S6 | Growth |
| Stromal Markers | | P-MTOR | Growth |
| Fibronectin | Matrix Glycoprotein | Nuclear | |
| SMA | Myoepithelial | Histone H3 | Chromatin |
| Vimentin | Mesenchymal | Cell Death | |
| Epigenetic Mark | | cleaved PARP | Apoptosis |
| H3K27 | Gene Repression | cleaved Casp3 | Apoptosis |

b

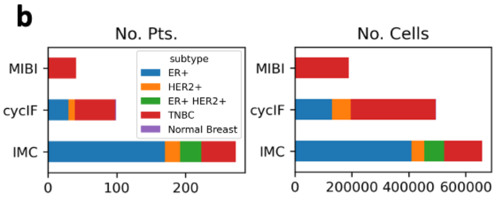


c

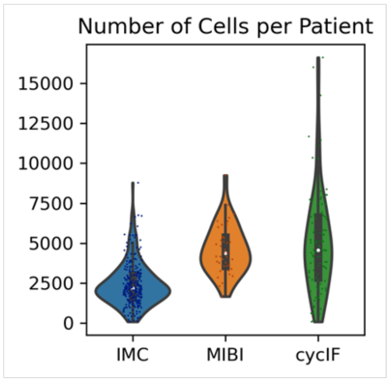


**S4. Comparison of breast cancer MXI panels and dataset sizes.**

a. The antibody panels from the three breast cancer datasets: our own CyCIF data from two breast cancer TMAs and publicly available IMC and MIBI data, including markers for cytokeratins, adhesion molecules, hormone receptors, receptor tyrosine kinase (RTK) signaling, cell growth and division, endothelial, immune and stromal cells. b. Number of patients (left) and cells (right) in each dataset, colored by subtype. c. Number of cells per patient in each dataset.

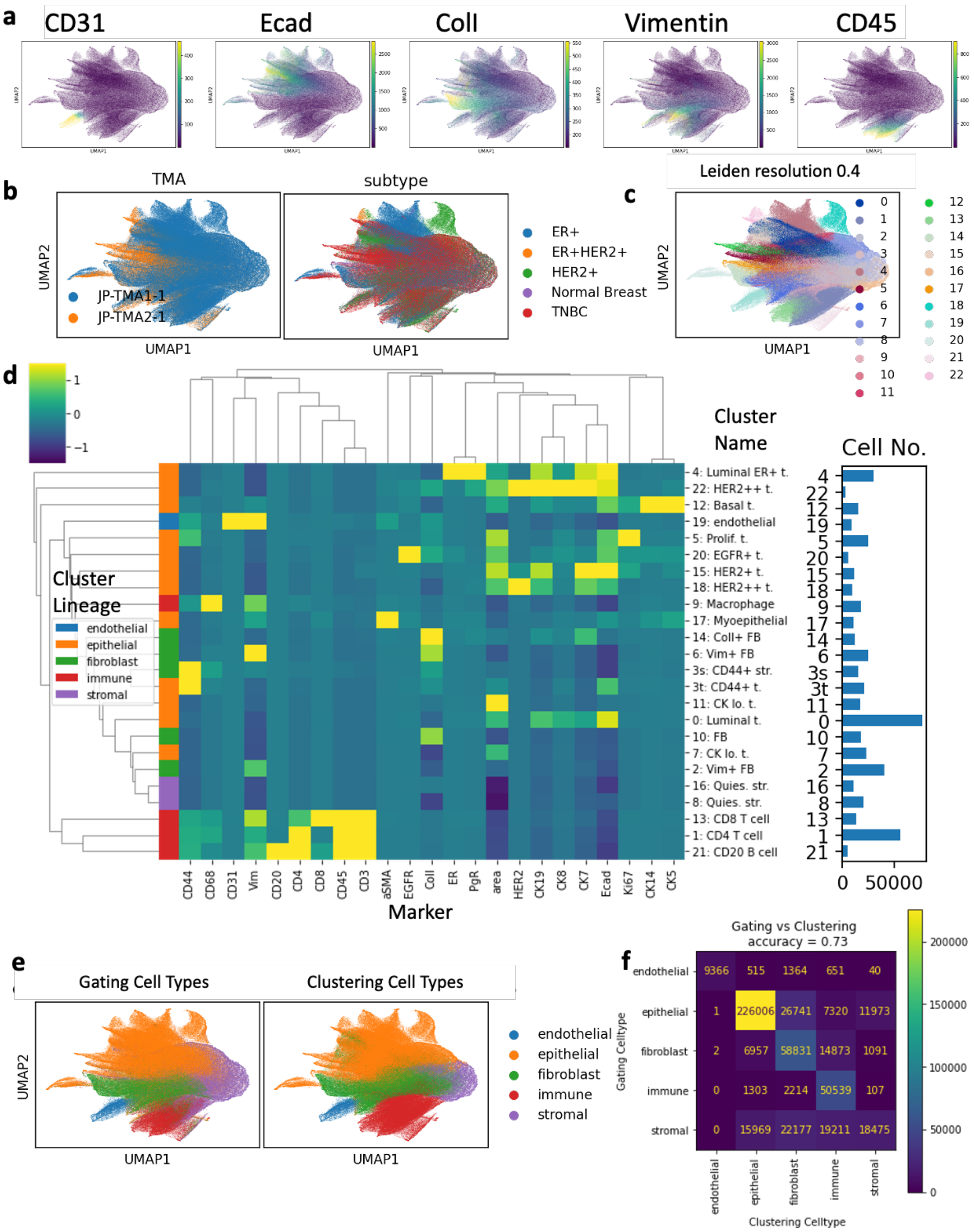


### S5. Single Cell Analysis of CyCIF data

a. Single-cell segmentation and feature extraction were done with mplexable. A UMAP embedding was generated based on single-cell mean intensity values (30 k-nearest neighbors). The UMAP is colored by cell lineage markers CD31, endothelial, E-cadherin (Ecad) epithelial, collagen I (ColI) and vimentin, fibroblast, and CD45, immune. b. UMAP colored by TMA (left) and breast cancer subtype (right). c. Unsupervised clustering with the Leiden algorithm (resolution 0.4) resulted in 23 cell types. d. Heatmap of mean fluorescence intensity of each marker in CyCIF cell type clusters. Twenty-two markers and one morphology feature (nuclear area) were used for clustering. Cell types were annotated as endothelial, epithelial, fibroblast, immune or stromal (left color bar on heatmap) and named based on marker expression (right labels on heatmap). e. Manual gating of the markers in (a) were used to determine cell types (left) separately from the Leiden annotated cell lineages (right). f. Confusion matrix of gating-based versus clustering-based cell lineages shows 73% accuracy.

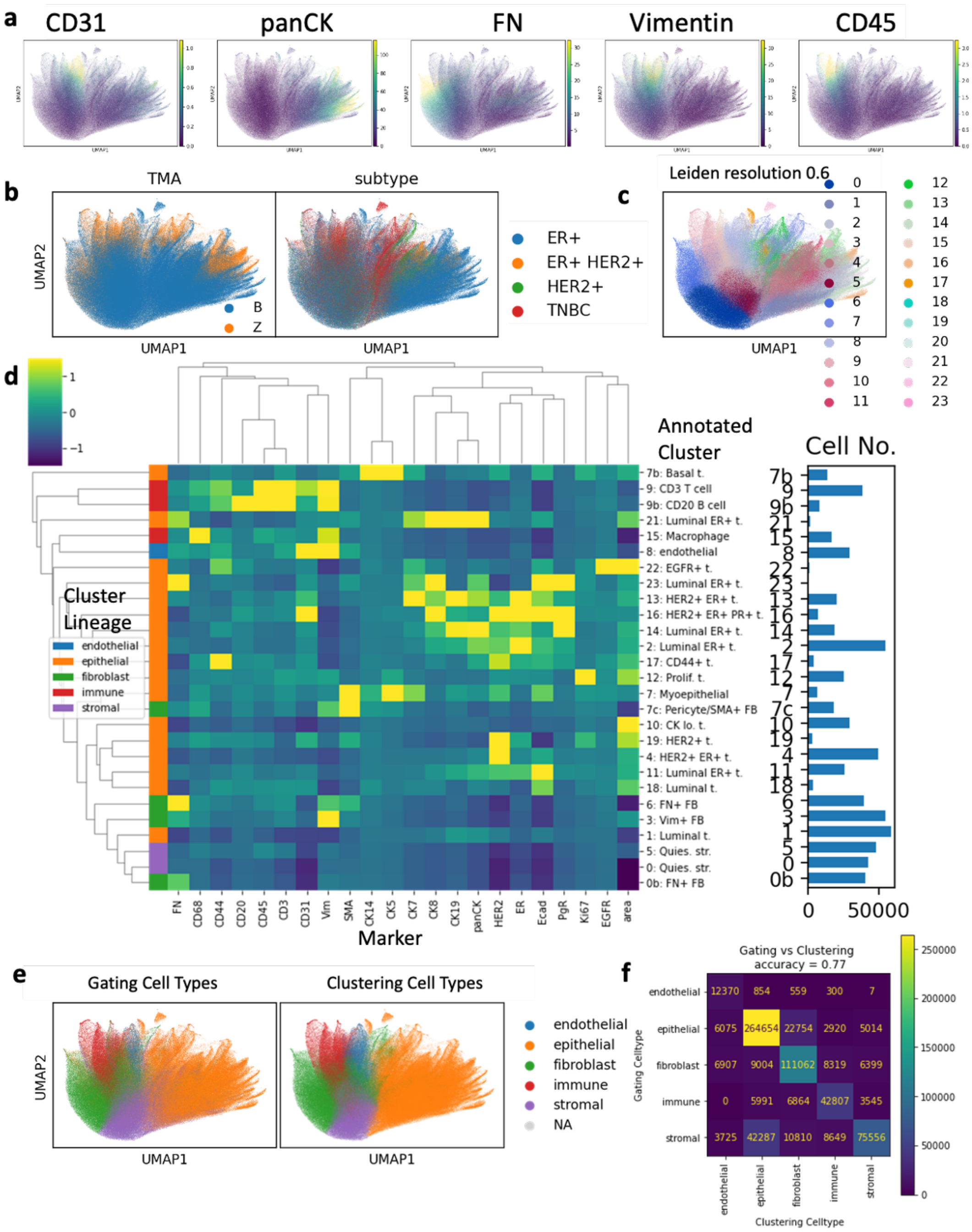


### S6. Single Cell Analysis of IMC data

a. Single-cell segmentation and feature extraction were done with mplexable. A UMAP embedding was generated based on single-cell mean intensity values (30 k-nearest neighbors). The UMAP is colored by cell lineage markers CD31, endothelial, pan-cytokeratin (panCK) epithelial, fibronectin (FN) and vimentin, fibroblast, and CD45, immune. b. Two TMAs (left) and four subtypes (right) were clustered together for cell typing. c. Unsupervised clustering with the Leiden algorithm (resolution 0.6) resulted in 25 cell types. d. Heatmap of mean fluorescence intensity of each marker in IMC cell type clusters. Twenty-one markers and one morphology feature (nuclear area) were used for clustering. Cell types were annotated as endothelial, epithelial, fibroblast, immune or stromal (left color bar) and named based on marker expression (right). e. Manual gating of the markers in (a) were used to determine cell types separately from the Leiden annotated cell types. f. Confusion matrix of gating-based versus clustering-based cell lineages shows a 77% agreement.

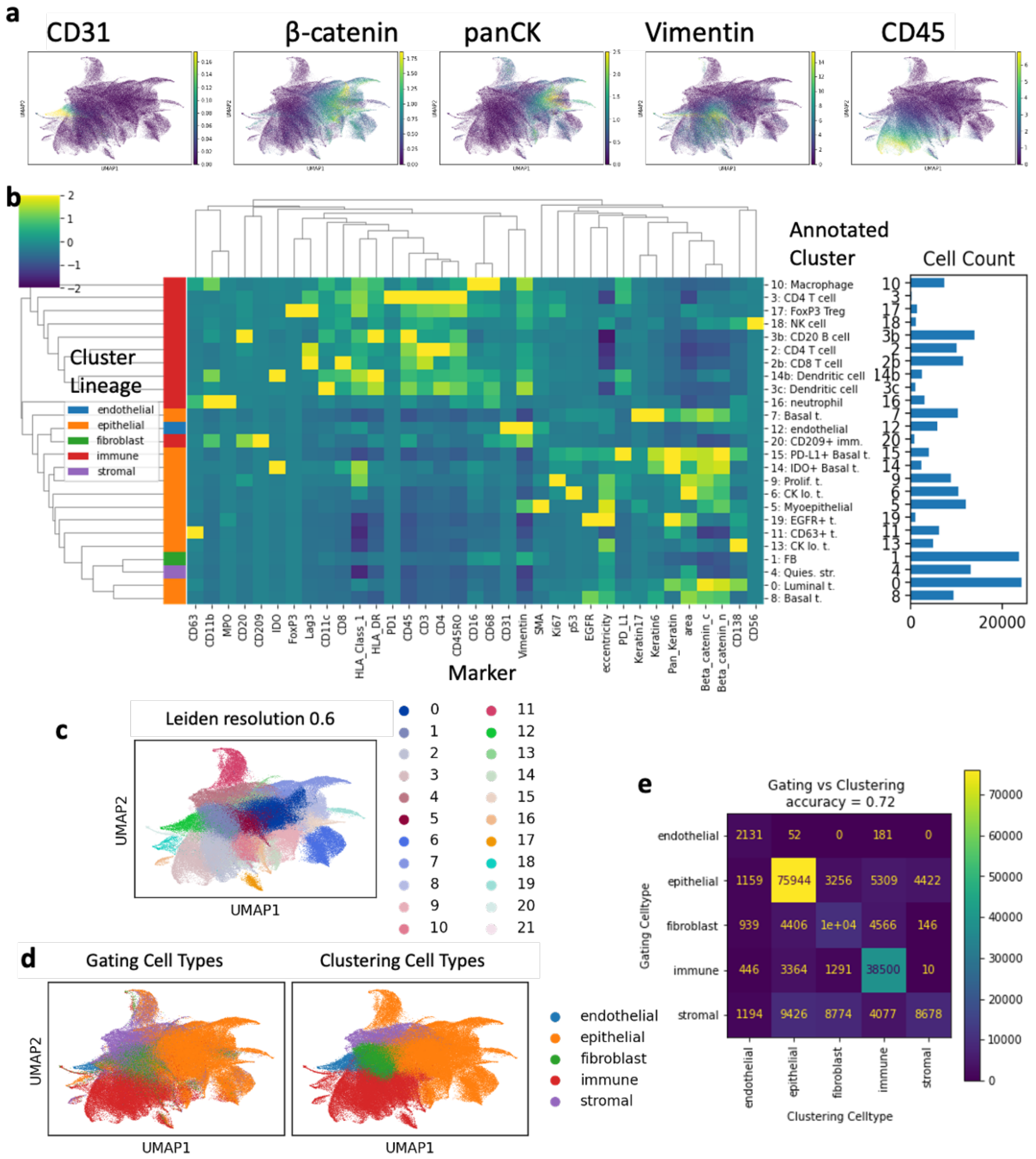


#### S7. Single Cell Analysis of MIBI data

a-e. Unsupervised clustering defines cell types in MIBI data. a. Single-cell segmentation and feature extraction were done with mplexable. A UMAP embedding was generated based on single-cell mean intensity values (30 k-nearest neighbors). The UMAP is colored by cell lineage markers CD31, endothelial, β-catenin and pan-cytokeratin (panCK) epithelial, vimentin, fibroblast, and CD45, immune. b. All samples were from a triple-negative breast cancer TMA. Thirty-three markers and one morphology feature (nuclear area) were used for clustering. Cell types were annotated as endothelial, epithelial, fibroblast, immune or stromal (left color bar) and named based on marker expression (right). c. Unsupervised clustering with the Leiden algorithm (resolution 0.6) resulted in 22 cell types. d. Manual gating of the markers in (a) were used to determine cell types separately from the Leiden annotated cell types. e. The gating-based versus clustering-based cell types had a 72% agreement.

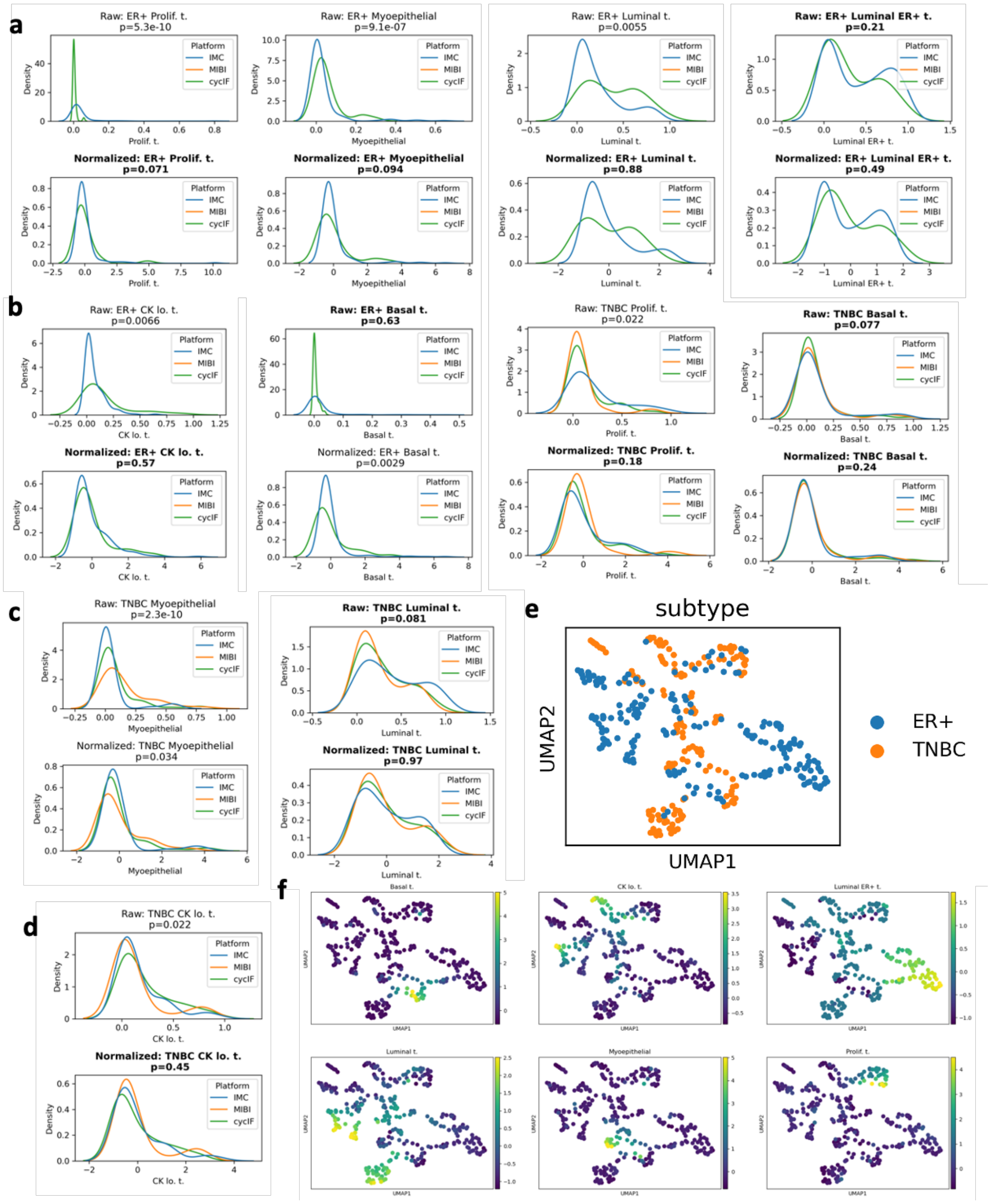


### S8. Normalization of epithelial fractions across platforms.

a-d. Kernel density estimates of fraction of epithelial cells of each phenotype before (top) and after (bottom) normalization. P-value in figure title is significant difference between platforms by Kruskal-Wallis H-test. Title text is bolded for p-values >0.05 indicating normalization resulted in no significant differences in median abundance of cell types between the platforms. e. UMAP embedding of patients by fraction of epithelial cell types in all tumor cells, colored by clinical subtype. f. UMAP embedding from (e), colored by marker abundance.

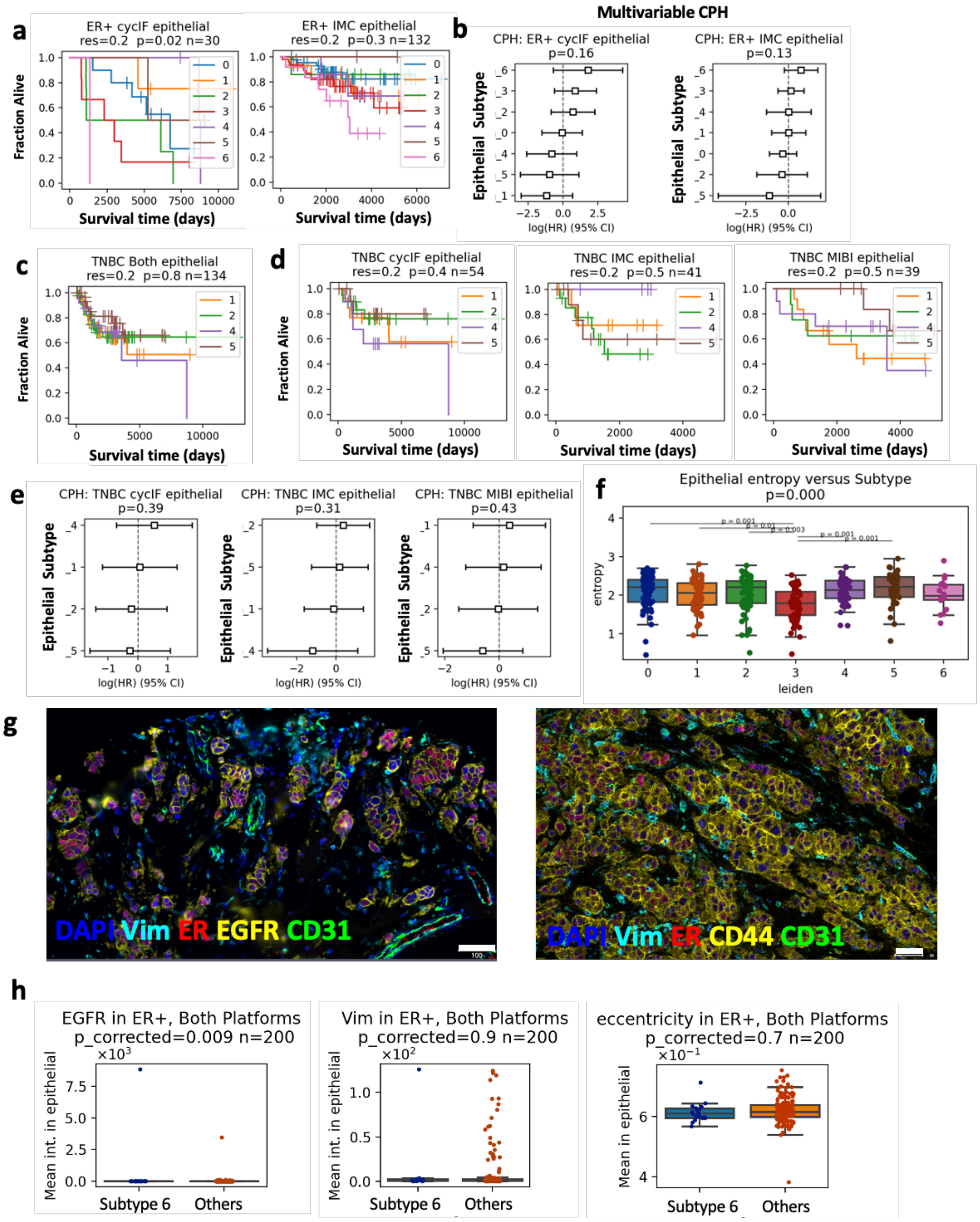


#### S9. Prognostic Value of Epithelial Subtypes.

a. Kaplan-Meier (K-M) curves (p-value from log-rank test) comparing overall survival (OS) in epithelial subtypes in ER+ tumors, by platform. b. Cox proportional hazard (CPH) modelling of epithelial subtypes versus overall survival in ER+, by platform. c. K-M OS curves of epithelial subtypes in all TNBC. d. K-M OS curves of epithelial subtypes in TNBC, by platform. d. CPH modelling of epithelial subtypes OS in TNBC, by platform. a-d. Platform, N number of patients and p-value shown in figure title. f. Shannon entropy of patients' epithelial phenotypes in each epithelial subtype. g. Example images of subtype 6 in ER+ tumors from the cycIF (left) and IMC (right) cohorts. scale bar = 100 microns. h. Mean intensity of selected markers and nuclear eccentricity of ER+ patients in poor-prognosis subtype 6 versus other ER+ patients. p-values obtained from t-tests and corrected for multiple testing with the Benjamini–Hochberg method.

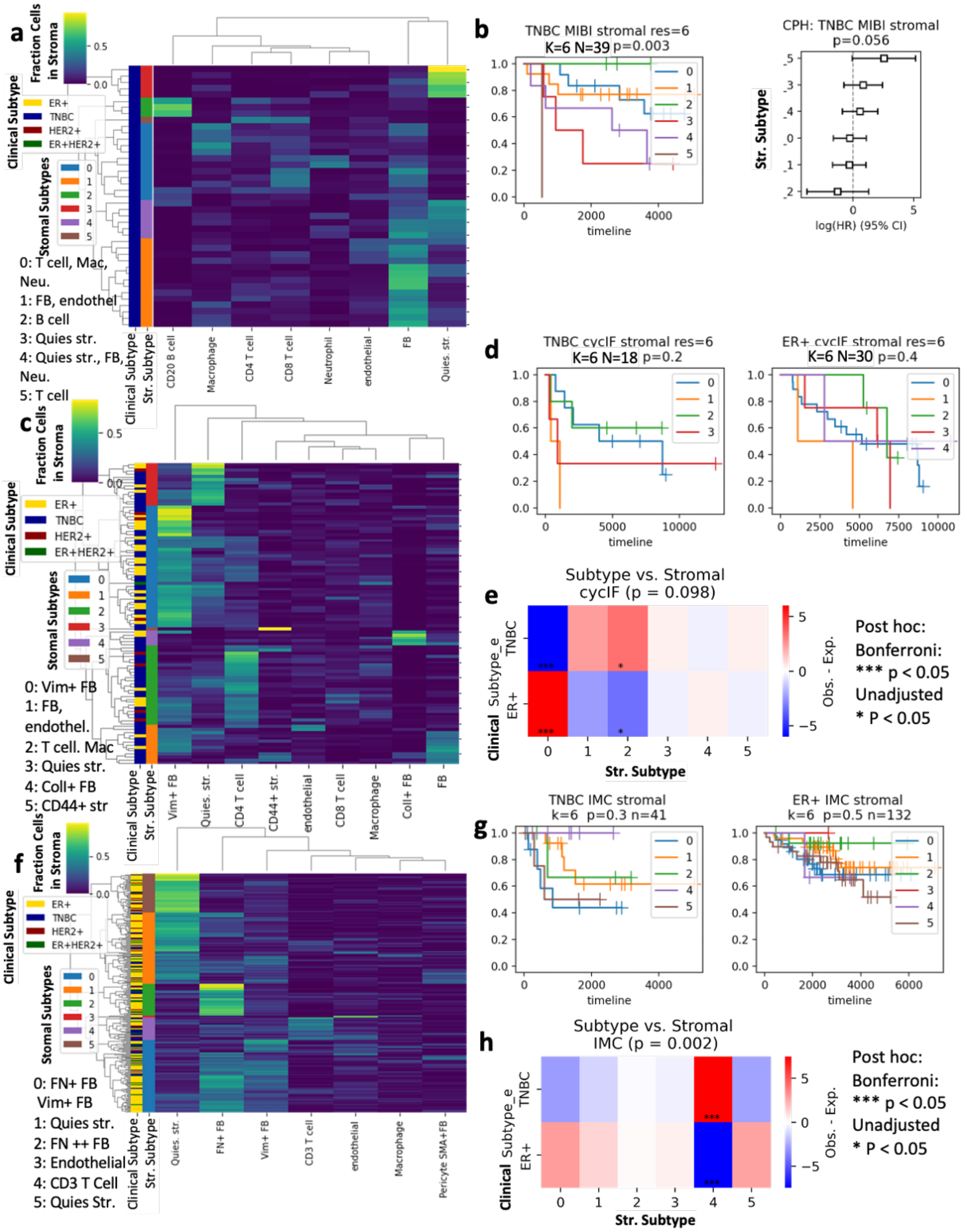


### S10. Prognostic Value and Clinical Subtype Correlation of Stromal Subtypes.

a, c, f. All patient tissues from each platform were hierarchically clustered based on the fraction of the common stromal cell types (>2%) in all stromal cells, selecting k=6 stromal subtypes. b. MIBI Kaplan-Meier curves (p-value from log-rank test) and Cox proportional hazard (CPH) models comparing OS in stromal subtypes, n=39 patients. c, d. CyCIF Kaplan-Meier curves comparing OS in stromal subtypes (p-value from log-rank test) n=18 TNBC, 30 ER+ patients. f, g. IMC Kaplan-Meier curves comparing OS in stromal subtypes (p-value from log-rank test) n=41 TNBC, 132 ER+ patients. e, h. Observed minus expected number of patients for each clinical subtype versus stromal subtype for CyCIF (e) and IMC (h). Overall p-value given in title (Chi-squared) and pairwise Bonferroni adjusted (***) and unadjusted (*) p-values < 0.05 marked on heatmap cells.

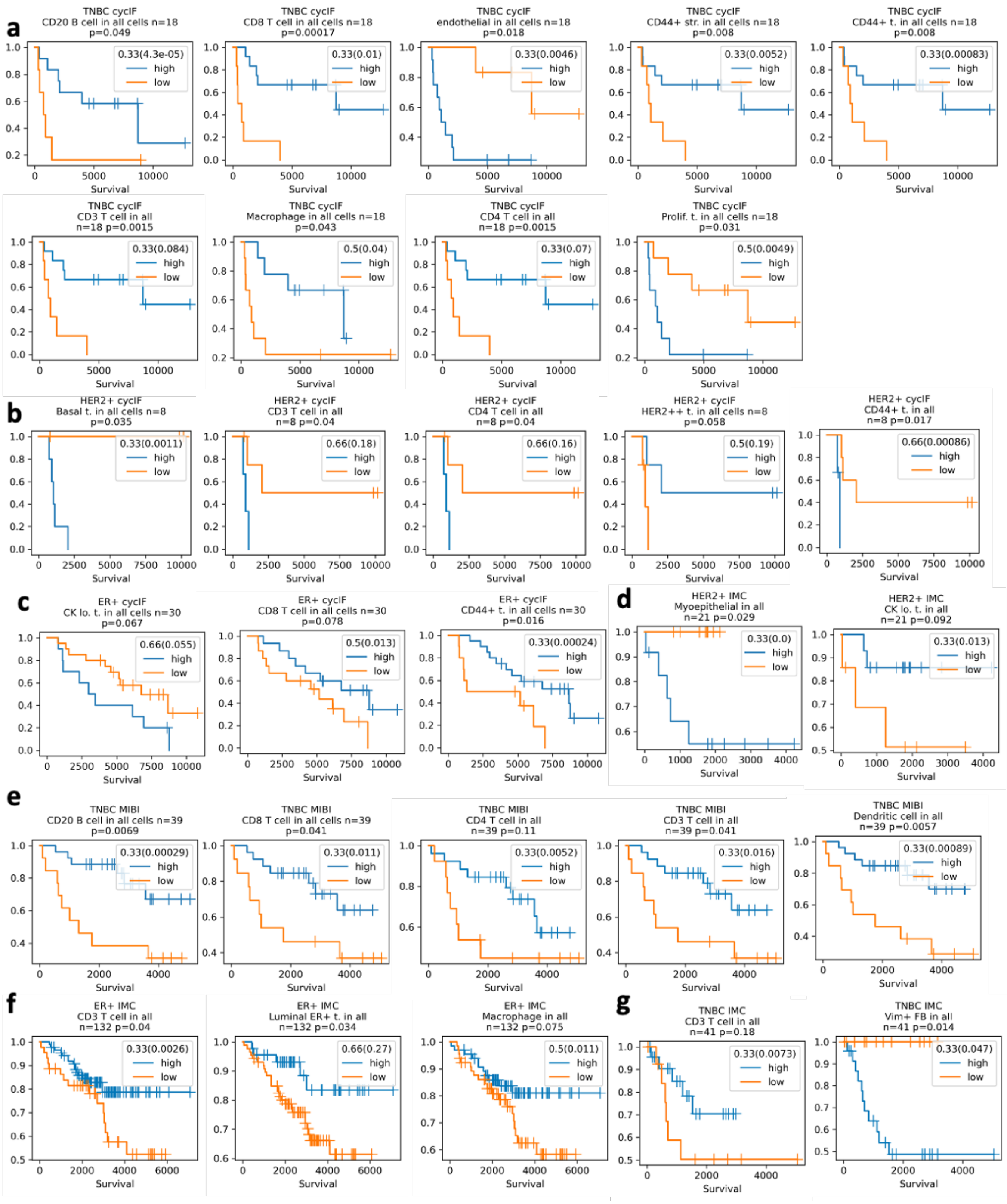


**S11. Single Variable Prognosis in separate cohorts.**

a-c . Kaplan-Meier (K-M) curves of high/low abundance of various cell types versus OS in CyCIF cohort for TNBC (a), HER2+ (b) and ER+ (c). d-g. Kaplan-Meier (K-M) curves of high/low abundance of cell types versus OS in IMC and MIBI cohorts. a-e. Cut-off values (tertiles of median) given in K-M legends. K-M p-values derived from log-rank test and given in figure titles, along with subtype and n number of patients.

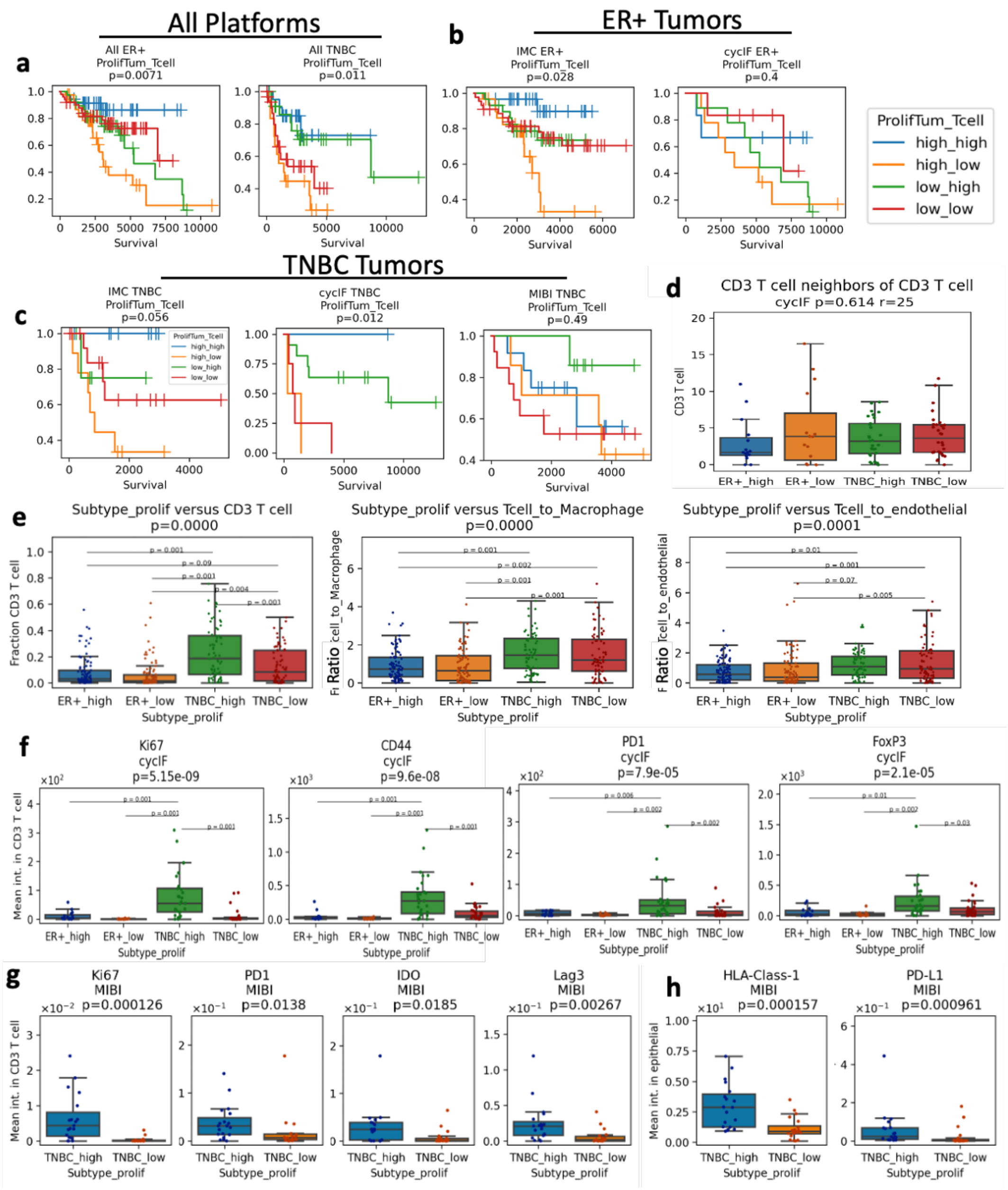


**S12. Prognostic value of proliferation and T cell abundance.**

a-c. Kaplan-Meier analysis of overall survival in each subtype split by proliferation low/high and CD3 T cell low/high, in combined cohorts (a), and from separate platforms' ER+ (b) and TNBC patients (c). p-values (log-rank) given in figure title. d. Mean number of T cell neighbors of T cells within a 25 µm radius in tissues from high and low proliferation ER+ or TNBC tumors in the CyCIF cohort. e. Mean fraction of T cells and ratio of T cells to macrophages and T cells to endothelial cells in tissues from high and low proliferation ER+ or TNBC tumors in the CyCIF and IMC cohorts. f. Ki67, CD44, PD1 and FoxP3 intensity in T cells indicating proliferation, memory/effector, checkpoint and regulatory function in tissues from high and low proliferation ER+ or TNBC tumors in CyCIF cohort. g. Ki67, PD1, IDO and Lag3 intensity in T cells indicating proliferation and checkpoint function in tissues from high and low proliferation TNBC tumors in MIBI cohort. h. HLA-Class-1 and PD-L1 in epithelial cells indicating antigen presentation and checkpoint in tissues from high and low proliferation TNBC tumors in MIBI cohort. d-f. Kruskal-Wallis H-test P-value given in figure title. Post-hoc Tukey HSD used for pairwise comparisons between groups. g-h. P-value from Mann-Whitney U rank test given in figure title.

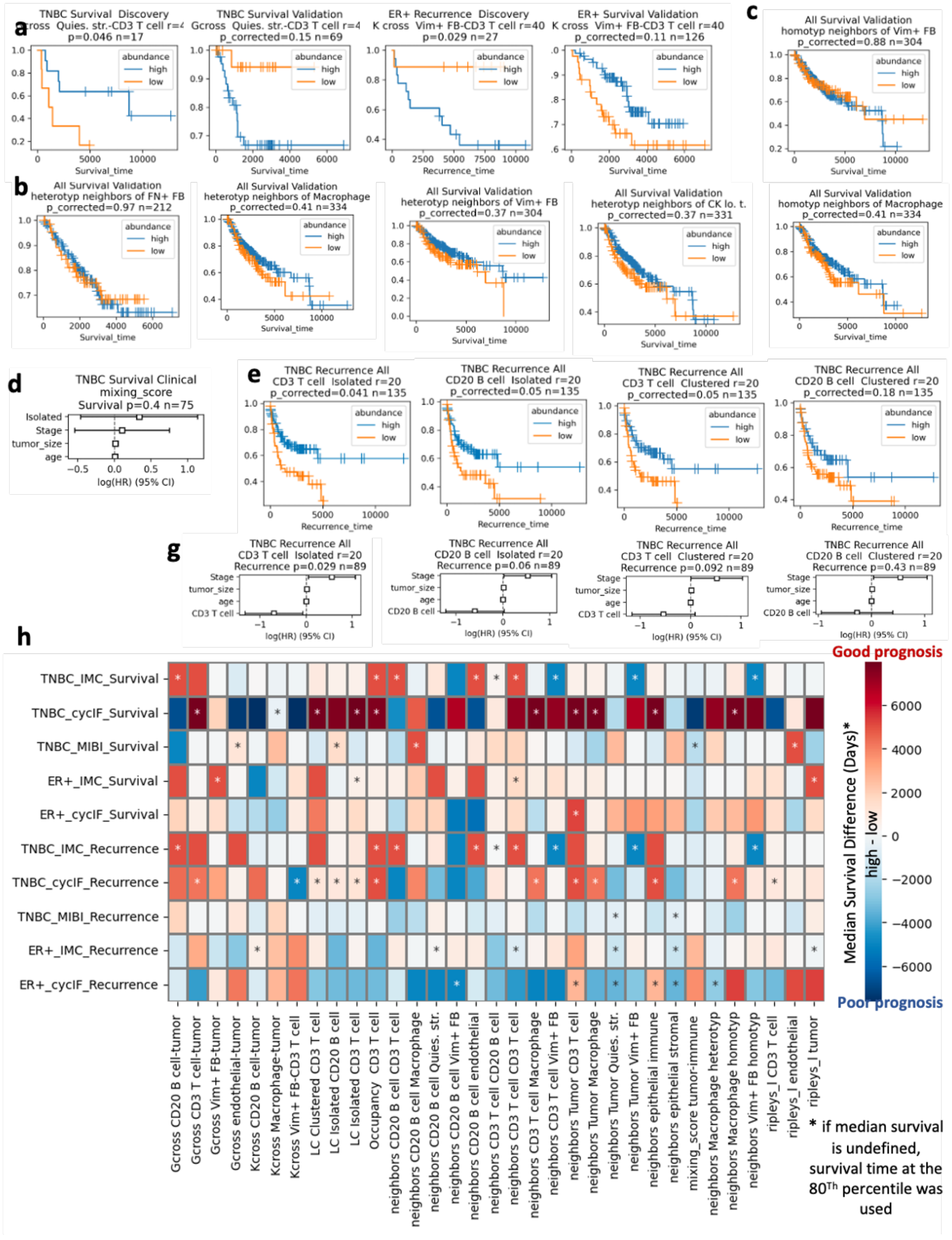

#### S13. Prognostic value of tumor immune spatial metrics

a. CPH modelling of mixing score versus overall and recurrence-free survival in TNBC. b. Prognostic value of CD3 T cell spatial aggregates in in TNBC MIBI cohort (OS, p-value from log-rank test) and CyCIF/IMC with clinical variables (RFS, p-value from CPH model). c.-e. Opposite survival associations in different cohorts for e. tumor to quiescent stroma Kcross function, 75 µm radius, d. macrophage neighbors of CD3 T cell, 25 µm radius, e. vimentin+ fibroblast neighbors of proliferating tumor, 25 µm radius. b-e. Kaplan-Meier survival curves. p-value from log-rank test and n number of patients given in figure title.

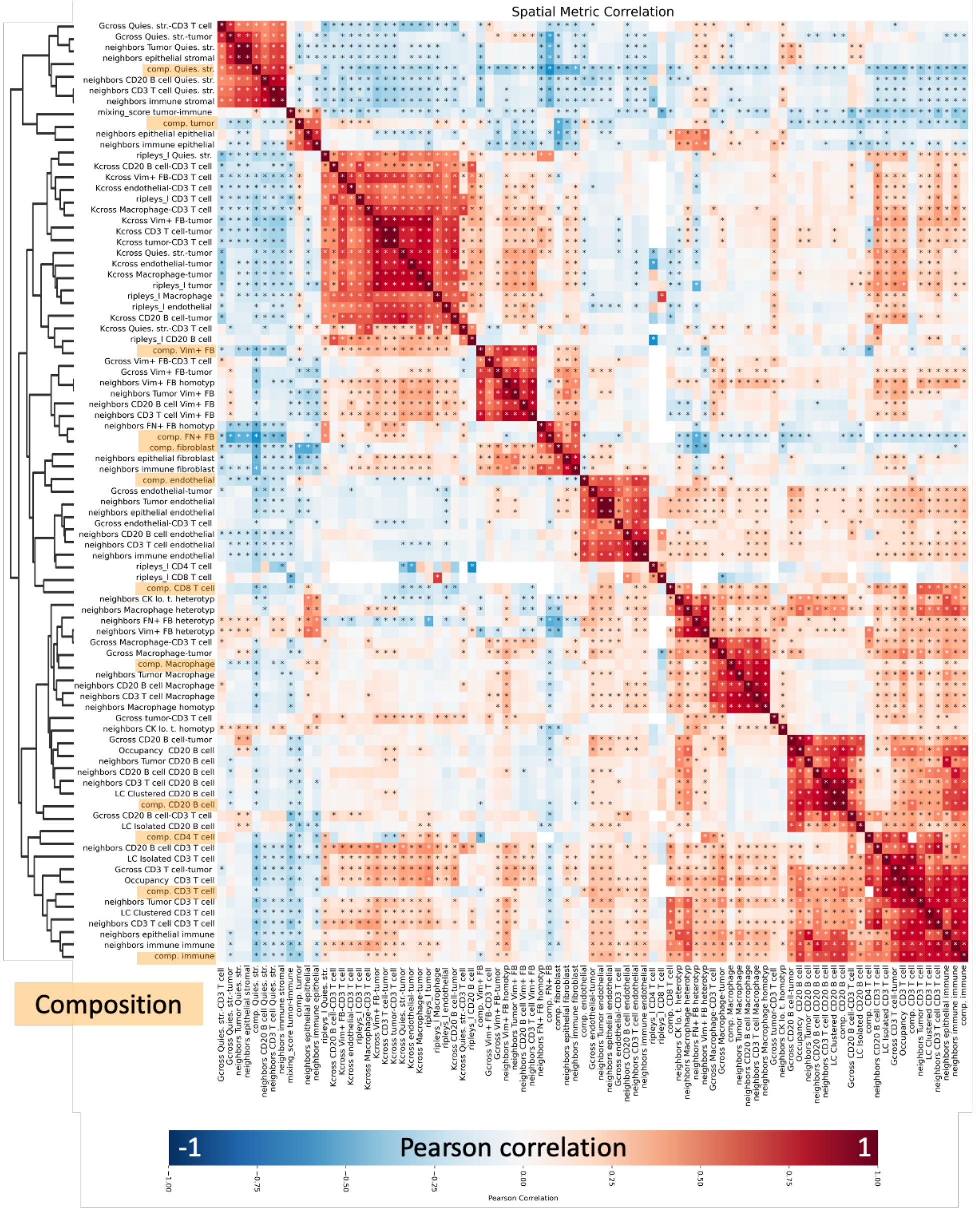


**S14. Correlation of Spatial Metrics and Tissue Composition.**

Heatmap of Pearson correlation between spatial metrics and tissue cell type composition (fraction of cells in tissue). Composition (comp.) variables are highlighted in orange. Asterisk denotes significant correlation ($p<0.05$). Dendrogram shows hierarchical clustering of metrics.

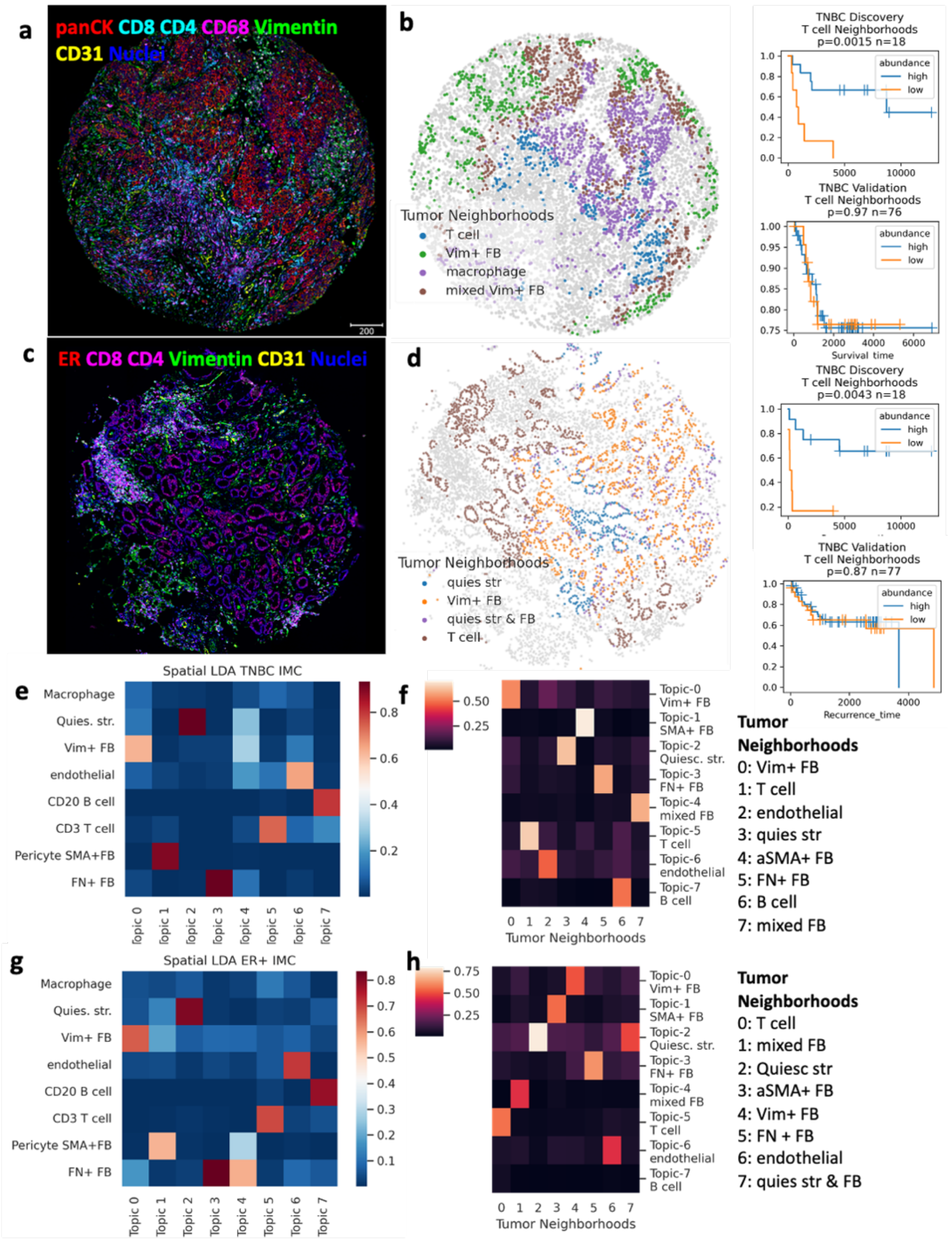


**S15. Spatial LDA Tumor Neighborhoods.**

a. CyCIF staining of tissue showing tumor (panCK), T cell (CD4 and CD8) fibroblast (vimentin), macrophage (CD68) and endothelial (CD31) markers. b. Tissue from (c) with tumor cells colored by their spattal LDA neighborhood cluster from (b). Tumor cells colored by T cell- (blue), macrophage- (purple), mixed fibroblast- (brown) and vimentin+ fibroblast-neighborhoods (green). c. CyCIF staining of tissue showing tumor (ER), T cell (CD4 and CD8) fibroblast (vimentin) and endothelial (CD31) markers. d. Tissue from (c) with tumor cells colored by their spatial LDA neighborhood cluster from (d). Tumor cells colored by T cell- (brown), quiescent stroma- (blue), mixed fibroblast- (purple) and vimentin+ fibroblast-neighborhoods (orange). e. Heatmap of stromal cell enrichment in spatial latent Dirichlet allocation (LDA) topic models of 100 µm tumor neighborhoods in TNBC tissue from the IMC platform. f. Heatmap of fraction of each topic in each neighborhood cluster resulting from K-means clustering (k=8) of spatial LDA topics from (a). g-h. heatmaps as defined in e and f, for ER+ tumors from the IMC cohort.

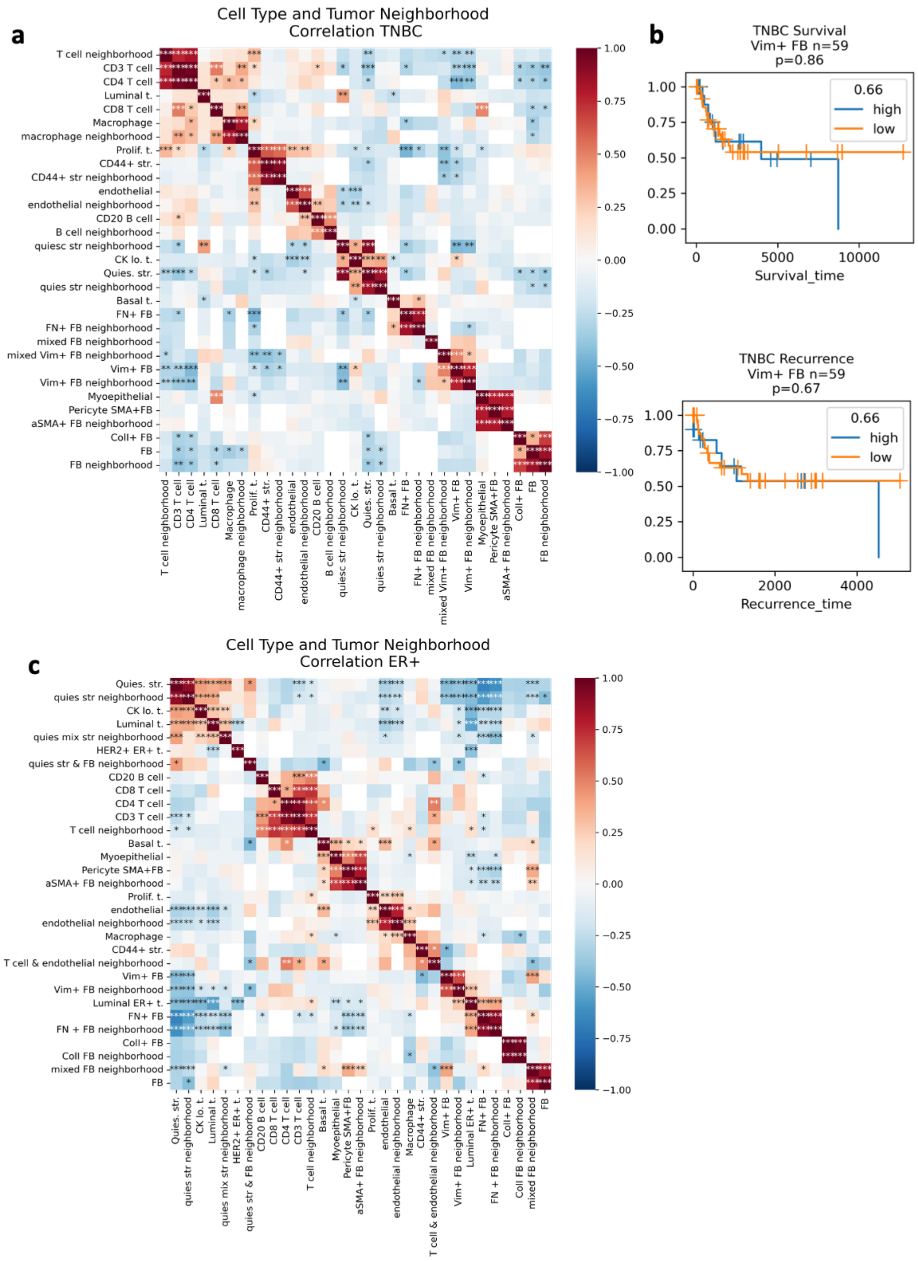


### S16. Correlation of tumor neighborhoods and tissue composition.

a. Heatmap of Pearson correlation between spatial LDA neighborhoods and fraction of cells in tissue in TNBC from combined CyCIF and IMC cohorts. Neighborhood variables are labelled as such and composition variables are just the cell type label. Asterisk denotes significant correlation ($p<0.05$). b. Kaplan-Meier OS and RFS curves of vimentin+ fibroblast abundance in TNBC tissues from combined CyCIF and IMC cohorts. c. Heatmap of Pearson correlation between spatial LDA neighborhoods and fraction of cells in tissue in ER+ tumors from combined CyCIF and IMC cohorts, labelled as in (a).

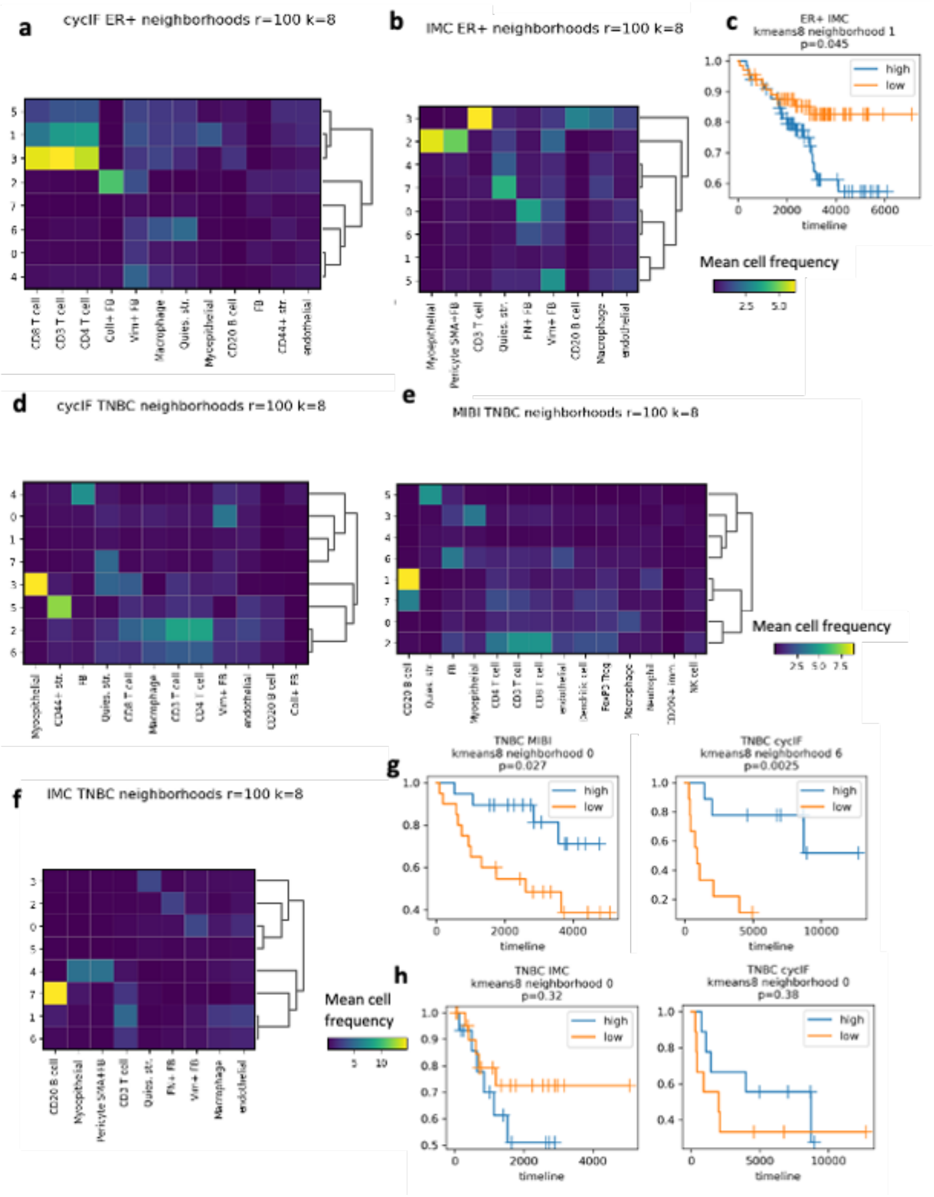


**S17. Neighborhoods defined by directly clustering the cell counts.**

a-b. Heat map of mean stromal cell frequency in neighborhood clusters in ER+ breast cancer tissues. Stromal cell counts in a 100 um radius surrounding epithelial cells were clustered with the kmeans algorithm, k=8, for the CyCIF (a) and IMC cohort (b). c. Kaplan-Meier analysis of a neighborhood from ER+ breast cancer (a-b) that was significantly associated with overall survival. d-f. Heat map of mean stromal cell frequency in neighborhood clusters in TNBC tissues from CyCIF (d), MIBI (e) and IMC cohort (f). g. Kaplan-Meier analysis of neighborhoods from TNBC (d-e) that were significantly associated with overall survival. h. Kaplan-Meier analysis of Vim+ FB neighborhoods in TNBC versus overall survival. c, g-h. High and low neighborhood frequency defined by the median of the cohort. p-values from log-rank test given in figure title.
